# Structure of Trypanosoma peroxisomal import complex unveils conformational dynamics

**DOI:** 10.1101/2023.04.03.535445

**Authors:** Ravi R. Sonani, Artur Blat, Malgorzata Jemiola-Rzeminska, Oskar Lipinski, Stuti N. Patel, Tabassum Sood, Grzegorz Dubin

**Affiliations:** Malopolska Centre of Biotechnology, Jagiellonian University, Krakow, Poland; Department of Biochemistry and Molecular Genetics, University of Virginia School of Medicine, Charlottesville, VA 22903, USA; Doctoral School of Exact and Natural Sciences, Jagiellonian University, Krakow, Poland; Faculty of Biochemistry, Biophysics and Biotechnology, Jagiellonian University, Krakow, Poland

## Abstract

Peroxisomes are membrane enclosed organelles hosting diverse metabolic processes in eukaryotic cells. Having no protein synthetic abilities, peroxisomes import all required enzymes from the cytosol through a peroxin (pex) import system. Peroxisome targeting sequence 1 (PTS1)-tagged cargo is recognized by soluble receptor, Pex5. The cargo-Pex5 complex docks at the peroxisomal membrane translocon, composed of Pex14 and Pex13, facilitating translocation into the peroxisomal lumen. Despite its significance, the structural basis of this process is poorly understood. Here, we present a cryo-electron microscopy (cryo-EM) structure of cargo-Pex5-Pex14 ternary complex from *Trypanosoma cruzi,* the causative agent of human Chagas disease. The atomic model of the Pex5 (residues 327-653) bound to malate dehydrogenase (MDH, cargo) tetramer and Pex14_NTD_ (residues 21-85) reveals dynamic motions and secondary interfaces. Specifically, we observe a swinging motion of Pex5 at the MDH-Pex5 interface. Additionally, the noncanonical contact surfaces are observed at the MDH-Pex5 and Pex5-Pex14 interfaces. Mutational analysis of the former interface indicates that it does not significantly enhance the affinity, underscoring the dynamic nature of cargo-Pex5 interactions. The latter interface constitutes an extended binding site of Pex14_NTD_ over Pex5. We discuss the implications of these findings for understanding peroxisomal transport mechanisms.

**HIGHLIGHTS:**

1. The structure of the peroxisomal import ternary complex (MDH-Pex5_TPR_-Pex14_NTD_) is determined using cryo-EM.
2. Pex5 interface with the cargo (MDH) is characterized by significant conformational dynamics.
3. The extended binding site of Pex14_NTD_ on Pex5 is revealed.

## INTRODUCTION

Peroxisomes compartmentalize a set of important metabolic processes in eukaryotic cells including catabolism of fatty acids and polyamines, biosynthesis of ether phospholipids, glyoxylate cycle in germinating seeds, glycolysis in *Trypanosoma* and many others^1,2^. Despite lacking genetic material and protein synthesis machinery, peroxisomes efficiently perform their functions by importing enzymes (referred to as cargo) from the cytoplasm^3^. The import process is orchestrated by a complex network of peroxins (Pex) proteins, numbering over a dozen, which recognize specific peroxisomal targeting signal (PTS) sequences present on folded cargo proteins in the cytoplasm. Subsequently, the cargo is translocated into the peroxisomal lumen, and the Pex system is recycled for subsequent rounds of cargo import^4^. Remarkably, the translocation process via the Pex system occurs without unfolding of the cargo proteins.

The initial step of peroxisomal transport involves recognition of a more common PTS1 signal by soluble receptor Pex5 in the cytoplasm. The less common PTS2 signal requires an additional receptor, Pex7. The PTS1 signal consists of variants of a Ser-Lys-Leu-COOH consensus sequence at the C-terminus of cargos^4^. Once bound by Pex5 in the cytosol, the cargo-Pex5 complex is subsequently translocated across the membrane by Pex13 and Pex14 assists in the process^5^, but the structural details of the interactions and the mechanism of translocation are still insufficiently understood.

The primary interaction between cargo and Pex5 is well-characterized through structures of isolated PTS1-peptide binding to the C-terminal globular domain of Pex5, which adopts the tetratricopeptide repeat (TPR) fold^6,7^. The PTS1 peptide burrows into the central cavity of Pex5 TPR domain^7^. Three existing crystal structures detailing the protein cargo-bound Pex5 TPR domain demonstrate secondary, PTS1-independent interactions between the cargo and Pex5, but the extent to which secondary interactions contribute to cargo-Pex5 affinity varies significantly and thus the role of secondary interactions is poorly understood^8–12^. This ambiguity underscores the need for additional structural investigations to elucidate the general nature and significance of the secondary interactions.

The process of translocating the Pex5-cargo complex across the peroxisomal membrane is subject to ongoing debate. What is established is that the import of the Pex5-cargo complex involves assistance from membrane-spanning receptors, Pex14 and Pex13^5^. It has been suggested that Pex5-cargo docking onto the membrane is assisted primarily via interaction with Pex14. Some studies propose that cargo-Pex5, in conjunction with Pex14 and Pex13, facilitate the formation of a dynamic pore, or engage a preformed Pex13 pore, enabling translocation of the cargo^13–15^. Pex14 was additionally implemented in cargo extraction from the translocation pore. The nature and dynamics of the pore and exact sequence of events require further investigation.

The interaction between Pex5 and Pex14 is primarily mediated through the recognition of multiple Wxxx(F/Y) motifs on Pex5 by the N-terminal globular domain of Pex14 (Pex14_NTD_)^16–19^. This canonical interaction has been extensively studied, typically through a system of short peptides containing the Pex5 Wxxx(F/Y) motif binding to Pex14_NTD_^20^. However, a comprehensive understanding of the Pex14 binding site on Pex5 is still elusive, as structural data of full-length Pex5 (or its domains including the Wxxx(F/Y) motif) bound to Pex14 are currently unavailable. Moreover, the cooperative role of multiple copies of Pex14 at the Wxxx(F/Y) motifs within Pex5 in binding and/or release of the cargo remains poorly characterized^16–19^.

In this study, we present the structure of the ternary complex involving cargo-Pex5-Pex14_NTD_ from *Trypanosoma cruzi*, the pathogen responsible for human Chagas disease^21^. Our structural analysis unravels the dynamic interactions of Pex5 with its cargo and secondary interfaces between cargo-Pex5 and Pex5-Pex14_NTD_ the role of which is substantiated by evaluation of structure guided mutants. Furthermore, our results show that the Pex14_NTD_ binding at Pex5 proximal Wxxx(F/Y) site is not enough for structural reorganization of the Pex5 NTD and cargo release *in vitro*.

## RESULTS AND DISCUSSION

### Reconstitution of cargo-Pex5-Pex14_NTD_ ternary complex

Specialized peroxisomes, known as glycosomes, import all glycolysis-associated enzymes (cargo) from cytosol through pex import system and compartmentalize glycolytic reactions in *Trypanosoma*. The pex import process is a validated drug target in Chagas disease^22–24^. After assessing various cargos (glycolytic enzymes) and Pex constructs (Supplementary Table 2), we chose glycosomal malate dehydrogenase (MDH, residues 1-323, UniProt Q4DRD8) as a cargo, and full length Pex5 (residues 1-666, GenBank PBJ69826.1) and N-terminal domain of Pex14 (Pex14_NTD_; residues 21-85, GenBank RNC55913.1) for *in vitro* reconstitution of the ternary complex. MDH is a crucial enzyme for glycolysis auxiliary pathway maintaining the pool of malate / oxaloacetate^25,26^. All three proteins were individually expressed in *Escherichia coli* and purified to homogeneity. The complex was reconstituted *in vitro* by mixing purified proteins and separating the complex from non-complexed components using size exclusion chromatography (SEC). The SEC profile of mixed components showed an additional peak at a higher molecular weight, corresponding to the ternary complex (Supplementary Figure 1). The presence of all three proteins in this peak was confirmed by the SDS-PAGE (Supplementary Figure 1).

### Cryo-EM reconstructions and heterogeneity of the ternary complex

Upon initial cryo-EM data analysis, the tetrameric arrangement of MDH was observed, yet significant variability in the MDH:Pex5 stoichiometry was noted (Figure 1). Avoiding chemical crosslinking, which could potentially homogenize the population, we instead extracted information on compositional heterogeneity from the available dataset. Classification efforts yielded 2D class averages representing diverse compositions of complexes, prominently featuring densities corresponding to complexes with one Pex5 molecule (MP1), two Pex5 molecules (MP2), and three Pex5 molecules (MP3) bound to the MDH tetramer (Figure 1A). Notably, the density corresponding to the third Pex5 molecule in MP3 was relatively weak (Figure 1A), and no complex with four Pex5 molecules was observed.

**Figure 1:**
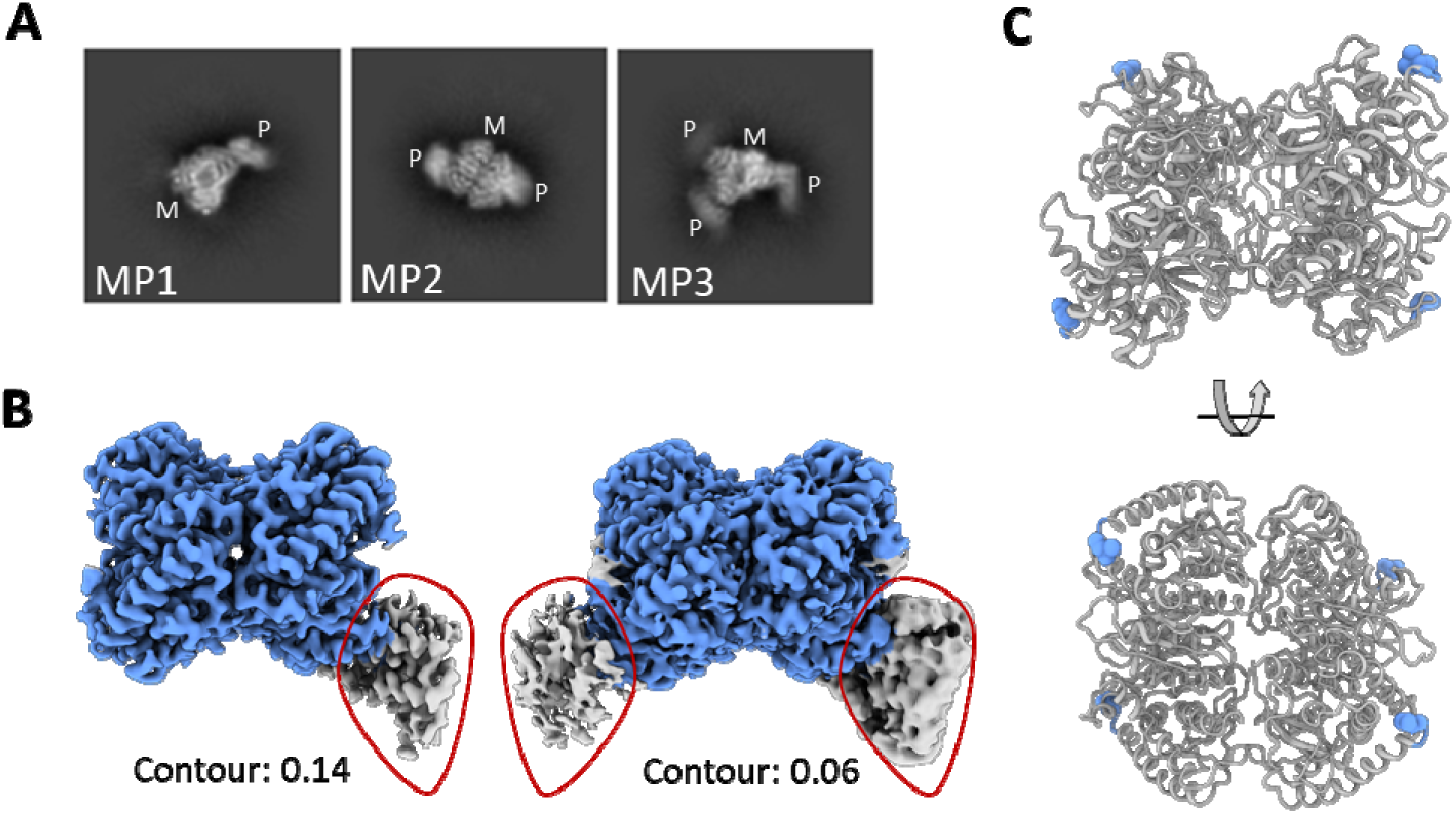
Cryo-EM analysis of peroxisomal import ternary complex – MDH-Pex5-Pex14_NTD_ through standard data processing pipeline. **(A)** 2D class averages showing density corresponding to the MDH-tetramer (M) and Pex5 (P). Three compositionally different complexes MP1, MP2 and MP3 were observed showing one, two and three copies of Pex5 bound to the MDH-tetramer, respectively. **(B)** 3D density maps of MP1 (left) and MP2 (right) complexes show clear density for MDH tetramer, but blurred density for the bound Pex5. Blurred density for Pex5 is encircled by red line. **(C)** The MDH tetramer structure shows the equal accessibility of all four PTS1 motifs (blue).

We attempted to reconstruct high resolution 3D maps of MP1, MP2 and MP3 using relevant particles by standard data processing procedures. While we could successfully reconstruct MP1 and MP2 maps showing one and two copies of Pex5 bound to MDH tetramer respectively (Figure 1B), the MP3 map showing density also for the third copy of Pex5 was not obtained most likely due to the low occupancy third copy and thus is not analyzed further. MP1 and MP2 maps show a well-resolved MDH region, but poorly resolved Pex5 regions, indicating conformational heterogeneity in the latter component (Figure 1). The resulting maps facilitated *de novo* model building for MDH, which forms a tetramer of approximately 10 nm in diameter. The peroxisomal targeting signal 1 (PTS1) motifs of all MDH subunits were found at the periphery of the tetramer and equally accessible to the solvent (Figure 1C).

To elucidate the conformational variability and improve the density of Pex region, we employed three-dimensional variability analysis (3DVA) using cryoSPARC. This approach improved density of Pex region and revealed significant motion of Pex5 relative to MDH, as depicted in Supplementary Movies 1 and 4, and facilitated the reconstruction of distinct 3D volumes representing snapshots of Pex5’s swinging motion over MDH within the MP1 and MP2 complexes. We refined and built two representative structures for each complex state, capturing two extreme conformations of the swinging motion namely, MP1-close (MP1-c), MP1-distal (MP1-d), MP2-close (MP2-c), and MP2-distal (MP2-d) (Figure 2; Supplementary Figure 2; Supplementary Movies 2, 3, 5, 6, and 7). The refined maps facilitated model building of MDH and the Pex5 TPR domain (residues 346-462, 487-653), including the upstream helix containing the Wxxx(F/Y) motif (residues 327-345), which had not been resolved in previously available Pex5 crystal structures^7^ (Supplementary Figures 3-6). However, the density corresponding to the Pex5 N-terminal domain (NTD; residues 1-326) was not discernible, indicating the flexible nature of this part of the molecule even in the presence of Pex14_NTD_, consistent with observations from prior studies^12,28^.

**Figure 2.**
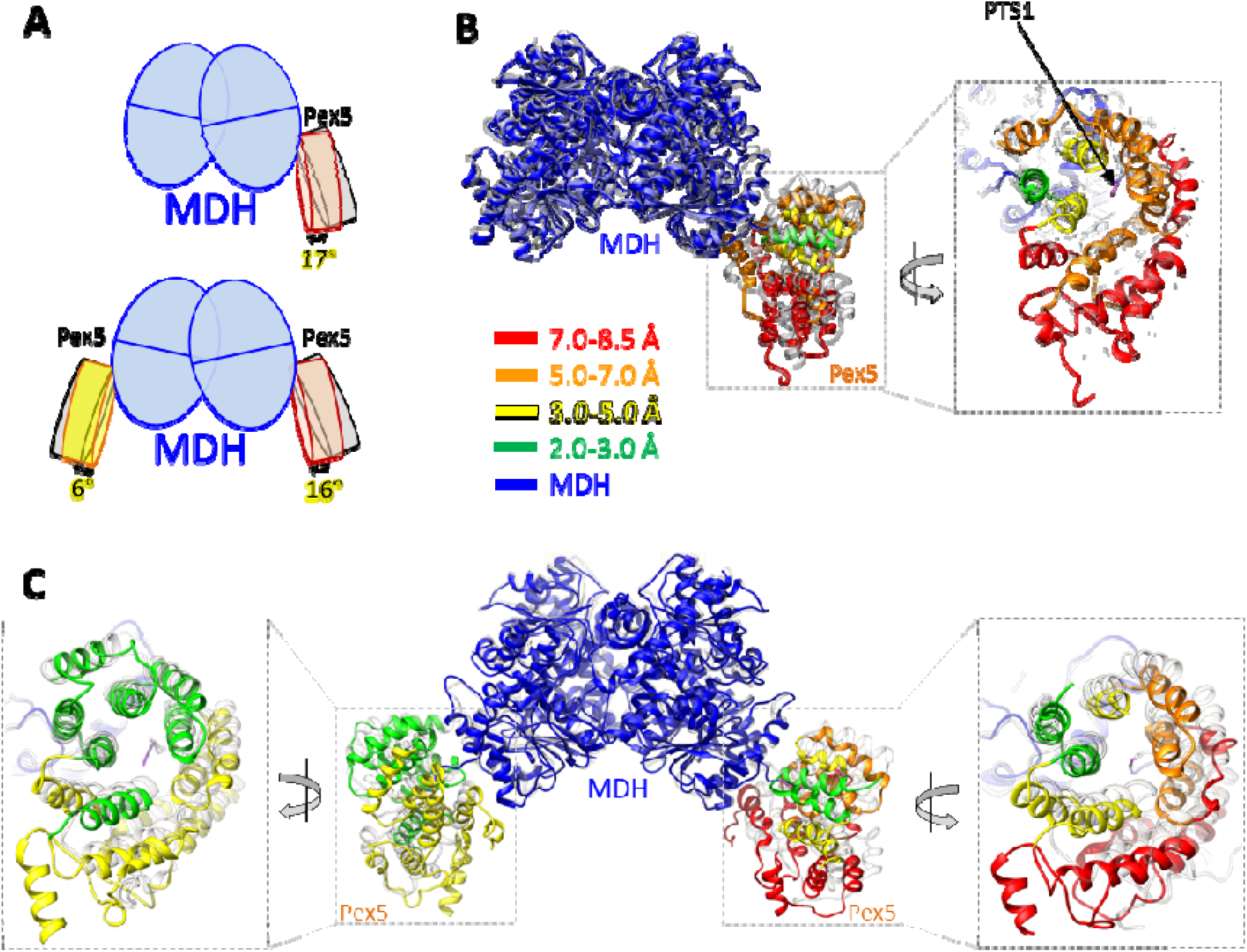
The Pex5-cargo interface shows significant swinging motions. **(A)** Schematic representation of angular movement (tilt) of Pex5 relative to the MDH in MP1 (upper panel) and MP2 (lower panel) structures. **(B, C)** Superimposed 3D structures of close (colored) and distal (grey) conformations of MP1 **(B)** and MP2 **(C)** complex. Color codes in proximal confirmation represent the extent of movement between distal and proximal confirmations as explained in inset. MDH is shown in blue on all panels. Pex5 is color coded according to the discussed features.

It is of note that although we observed particles with two Pex5 molecules bound on the same side of the MDH tetramer, we did not observe particles with two Pex5 molecules bound to diagonally opposite MDH subunits (Figure 2B). The underlying reason for this specific topology remains unclear as the PTS1 binding sites on all protomers are equally accessible (Figure 1C). Furthermore, the presence of particles representing the MP3 complex (Figure 1A), where three Pex5 molecules are bound to MDH tetramer, suggests there would be no steric clashes between Pex5 if diagonally bound. It is uncertain whether occupancy of more than one Pex5 site is necessary for translocation. The piggybacking of cargo isoforms within heteromultimeric complexes suggests that saturation of all Pex5 interaction sites is not required for translocation^29,30^. Furthermore, recent findings indicate that *in vivo*, only a single Pex5 molecule binds to a hexamer of cargo, implying that not all PTS1 sites need to be occupied by Pex5 for efficient translocation^12^.

### Pex5-binding on MDH is of dynamic nature

Recognition of the MDH by Pex5 is mediated via canonical interactions of PTS1 motif at the central cavity of Pex5 TPR domain (Figure 2B). This interaction is stabilized by a network of hydrogen bonds involving the PTS1 C-terminal carboxylate group, mainchain donors and acceptors, and lysine sidechain amine, as observed previously in different species^7^. The overall geometry of PTS1 bound Pex5 in *T. cruzi* is compact compared to the structure of apo-Pex5. Similar compaction is also observed for *Trypanosoma brucei* and human Pex5^6–8,31^, suggesting that TPR domain compaction upon PTS1 binding is general across species.

The 3D structural variability analysis reveals that Pex5, when bound to PTS1, exhibits swinging movements on the surface of MDH. We categorized the raw particles into 20 clusters, each representing varying orientation within this swinging motion covering the entire range of orientations between “distal” and “closed” conformations. The random distribution of particles among the clusters indicates a continuous spectrum of Pex5 swinging motion relative to MDH with no preferential orientations (Supplementary Figure 2). This swinging motion can be characterized by two components: tilting and twisting. Pex5, tilts nearly as a rigid body, oscillating between close and distal conformations with a maximum tilt angle of 17° relative to the PTS1 axis (Figure 2A; Supplementary Movies 1-7). Additionally, during this tilting motion, Pex5 twists up to 13° relative to the axis roughly parallel to the TPR3 motif (Supplementary Movie 8). The combined tilting and twisting motions result in a radial displacement of Pex5 ranging between 3 to 8 Å (Figure 2). Specifically, TPR motifs 1, 2, 6 and 7 (residues 330-425, 527-640) show displacements of 5-8 Å, which is greater than the central TPR motifs 3, 4 and 5 (residues 428-525), with displacements around 3.5 Å (Figure 2B, 2C; Supplementary Movies 1-8). 3DVA results further suggest that there is no coordination between the movement of two Pex5 molecules in MP2 complex (Supplementary Figure 8).

### Secondary MDH-Pex5 interfaces does not significantly contribute to affinity

Along PTS1 mediated interaction, two additional interfaces between MDH and Pex5 are identified by PISA analysis (Figure 3A). Interface 1 involves MDH residues 62-70 and Pex5 residues 439-444 forming a loop connecting α6 and α7. Notably, MDH residues 62-70 form a loop (MDH_62-70_) unique to the glycosomal isoform and absent in the cytosolic/mitochondrial isoforms and homologues from other organisms (Supplementary Figure 10), suggesting its possible specific role in glycosomal transport^32^. Interface 2 involves MDH residues 142-145 and Pex5 helix α15 (residues, 625-633) (Figure 3A).

**Figure 3:**
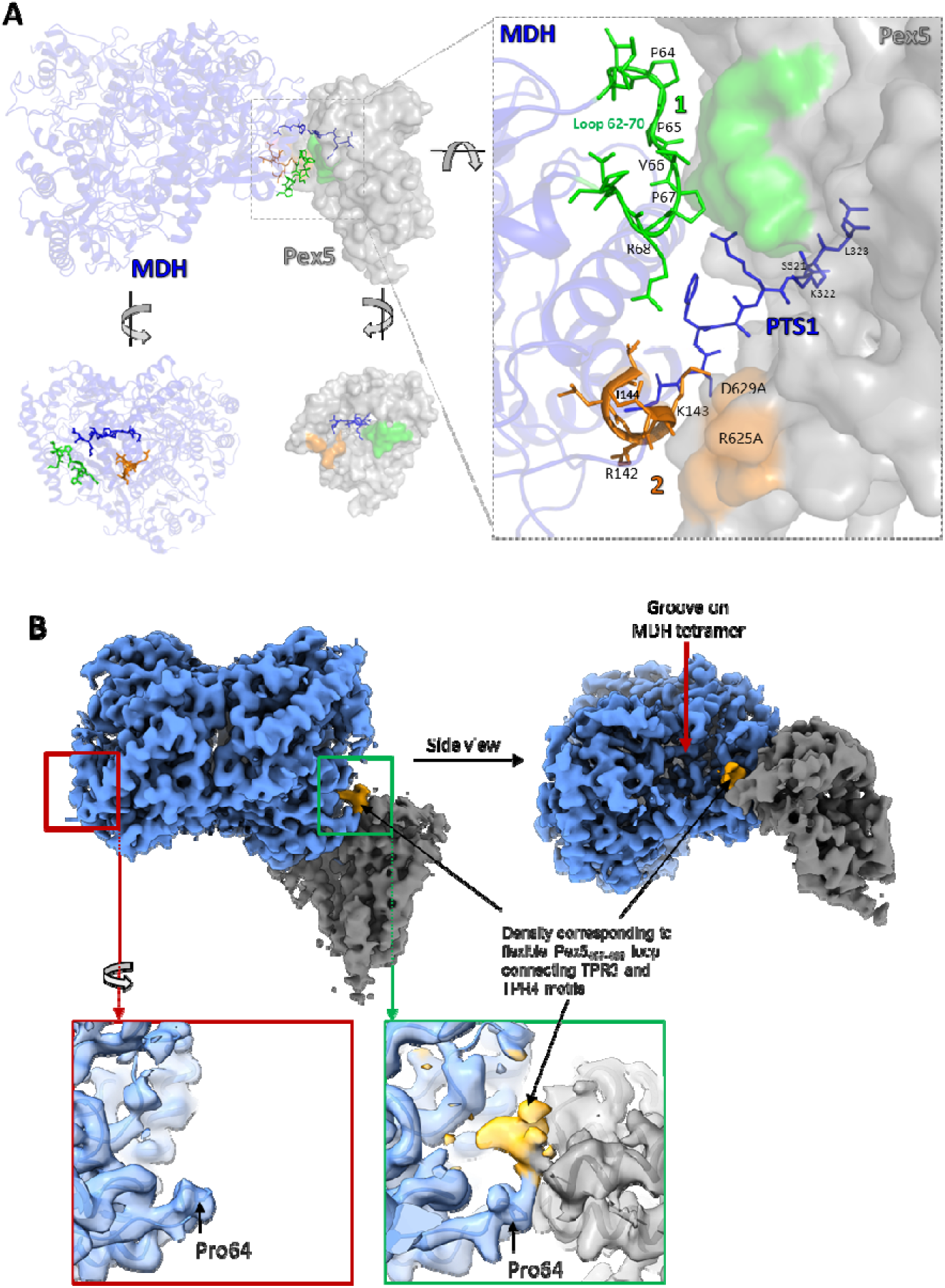
Non-canonical interactions of Pex5 and MDH. **(A)** Non-canonical interactions are represented by green and orange colours on MP1-d structure (upper panel), MDH only (lower left panel) and Pex5 only (lower right panel). MDH and Pex5 is depicted in blue ribbon and grey surface, respectively. Canonical PTS1 burrowing inside TPR-domain cavity is presented by blue-stick model. Right panel shows the enlarged view of MDH-Pex5 canonical PTS1 interface and noncanonical interactions. Density corresponding to the Pex5 loop transiently interacting with MDH_457-489_ groove in MP1-d map (gold). The density is not seen in a groove distal form Pex5 occupied site (red). See Supplementary Figure 12 for analogous analysis of MP2-d map.

Because close analysis of interfaces 1 and 2 does not reveal classical features of protein-protein interaction and because identified swinging motion affects the extent of the interfaces, we wanted to test if the identified interfaces contribute to affinity. Mutants were generated targeting residues at the MDH-Pex5 interface, and their affinities were determined by ITC (Table 1). The affinity of wild type Pex5 and MDH was estimated at K_d_ ∼ 435 nM (Table 1). To disrupt interface 1 we replaced the MDH_62-70_ loop with GS-linker. This resulted in a minor decrease (∼2.5-fold) in affinity towards wild type Pex5 indicating small contribution of the interface to affinity. Interface 1 at the Pex5 side is composed of the backbone atoms and thus not directly amenable to mutational analysis. To assess the contribution of interface 2, a double mutant of Pex5 (R625A and D629A) was constructed. The mutant exhibited an affinity comparable to the wild type Pex5, indicating insignificant contribution of interface 2 in MDH-Pex5 affinity. MDH side of the interface was composed of backbone atoms and thus was not analyzed by mutagenesis. To further explore the collective impact of secondary MDH-Pex5 interfaces, a linker (GSGS) was introduced between PTS1 and MDH to physically separate the binding partners thus disrupting any secondary interactions. The GSGS-linker mutant demonstrated nearly four times higher affinity compared to the wild type MDH (Table 1). The change in affinity was entropy driven indicating that the secondary interactions have overall negative effect on affinity by restricting the conformational dynamics of the wild type interface. The affinity of GSGS-linker mutant is comparable to that of a short peptide encompassing PTS1 of MDH (residues 319-ARSKL-323) (Table 1) indicating that the introduction of GSGS linker indeed abolished all secondary interactions at the Pex5-MDH interface.

**Table 1.**
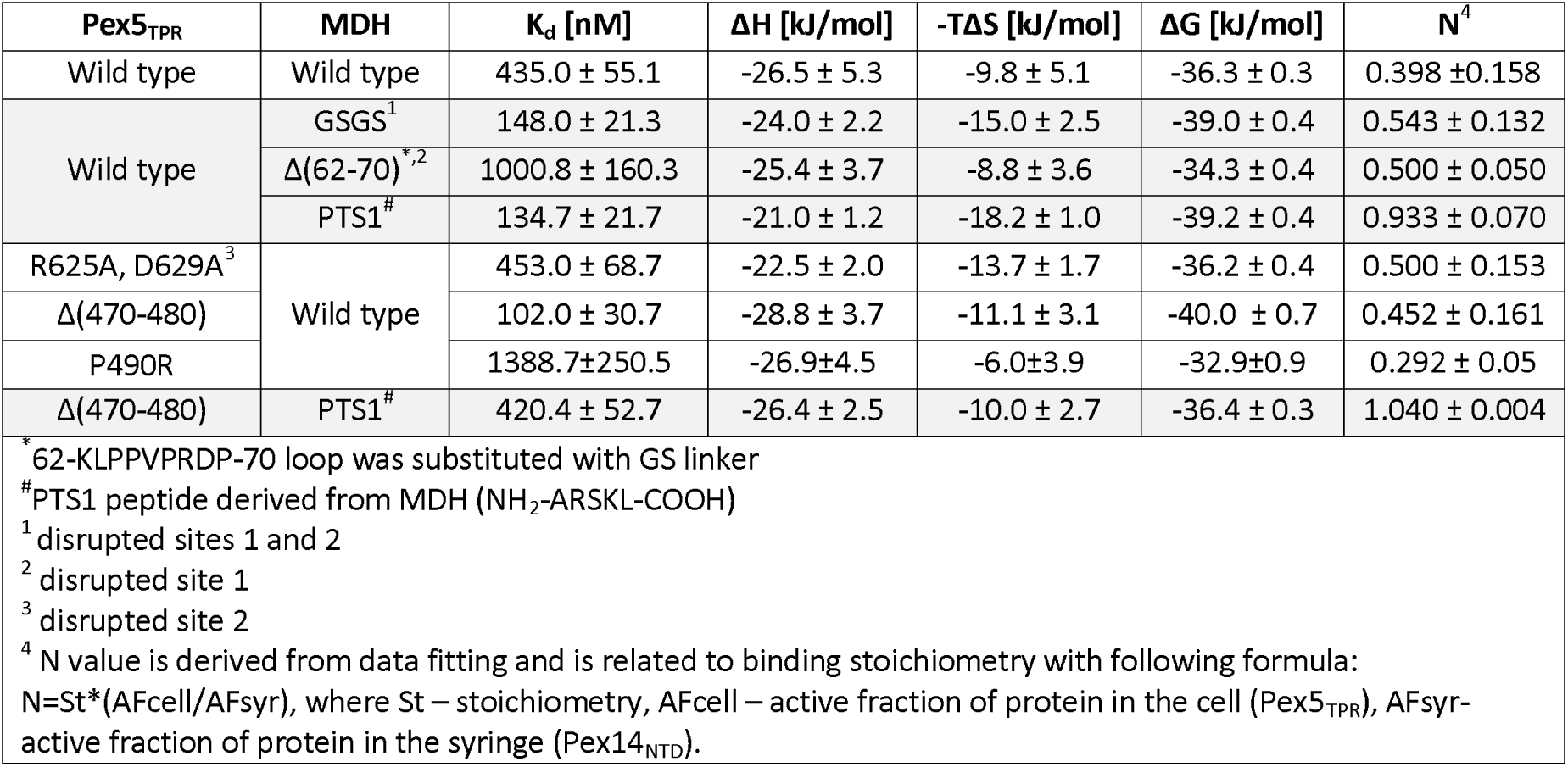
Mutational analysis of Pex5-cargo interface. Interactions of indicated mutants were characterized by Isothermal Titration Calorimetry (ITC).

| Pex5<sub>TPR</sub> | MDH | K<sub>d</sub> [nM] | ΔH [kJ/mol] | -TΔS [kJ/mol] | ΔG [kJ/mol] | N<sup>4</sup> |
| --- | --- | --- | --- | --- | --- | --- |
| Wild type | Wild type | 435.0 ± 55.1 | -26.5 ± 5.3 | -9.8 ± 5.1 | -36.3 ± 0.3 | 0.398 ± 0.158 |
| Wild type | GSGS<sup>1</sup> | 148.0 ± 21.3 | -24.0 ± 2.2 | -15.0 ± 2.5 | -39.0 ± 0.4 | 0.543 ± 0.132 |
|  | Δ(62-70)<sup>*,2</sup> | 1000.8 ± 160.3 | -25.4 ± 3.7 | -8.8 ± 3.6 | -34.3 ± 0.4 | 0.500 ± 0.050 |
|  | PTS1<sup>#</sup> | 134.7 ± 21.7 | -21.0 ± 1.2 | -18.2 ± 1.0 | -39.2 ± 0.4 | 0.933 ± 0.070 |
| R625A, D629A<sup>3</sup> | Wild type | 453.0 ± 68.7 | -22.5 ± 2.0 | -13.7 ± 1.7 | -36.2 ± 0.4 | 0.500 ± 0.153 |
| Δ(470-480) |  | 102.0 ± 30.7 | -28.8 ± 3.7 | -11.1 ± 3.1 | -40.0 ± 0.7 | 0.452 ± 0.161 |
| P490R |  | 1388.7 ± 250.5 | -26.9 ± 4.5 | -6.0 ± 3.9 | -32.9 ± 0.9 | 0.292 ± 0.05 |
| Δ(470-480) | PTS1<sup>#</sup> | 420.4 ± 52.7 | -26.4 ± 2.5 | -10.0 ± 2.7 | -36.4 ± 0.3 | 1.040 ± 0.004 |

<sup>\*</sup> 62-KLPPVPRDP-70 loop was substituted with GS linker <sup>#</sup> PTS1 peptide derived from MDH (NH<sub>2</sub>-ARSKL-COOH) <sup>1</sup> disrupted sites 1 and 2 <sup>2</sup> disrupted site 1 <sup>3</sup> disrupted site 2 <sup>4</sup> N value is derived from data fitting and is related to binding stoichiometry with following formula: $N = St * (AF_{cell} / AF_{syr})$ , where St – stoichiometry, AF<sub>cell</sub> – active fraction of protein in the cell (Pex5<sub>TPR</sub>), AF<sub>syr</sub> – active fraction of protein in the syringe (Pex14<sub>NTD</sub>).

The above findings collectively demonstrate that neither of the identified secondary interfaces significantly contributes to the MDH-Pex5 affinity, thus facilitating the restricted swinging motion of Pex5.

### Transient interaction of Pex5_457-489_ loop with MDH

The maps accounting for the distal states of MDH-Pex5 complex (MP1-d and MP2-d) show additional density proceeding Pex5 residue Ser491 and facing MDH_62-70_ loop (Figure 3B; Supplementary Figure 12). Notably, the Pex5_457-489_ loop (linking TPR motifs 3 and 4) is unresolved in all existing crystal structures, indicating its flexible nature. The density is present only at sites occupied by Pex5 and not empty PTS1 sites indicating it corresponds to a fragment of Pex5_457-489_ loop (Figure 3B; Supplementary Figure 12). Additionally, the density was absent in the maps of MDH tetramer solved in this study (Supplementary Figure 12) further suggesting it originates from Pex5. These results indicate restriction of conformational flexibility of Pex5_457-489_ loop in distal state of MDH-Pex5 complex.

The identified density was insufficient to construct an atomic model of the Pex5_457-489_ loop but indicates that loop extends into the groove between MDH subunits (Figure 3B). In glycosomal MDH, the groove is notably deeper (volume ∼4900 Å^3^) compared to non-glycosomal MDH (PDB id: 5ZI4) (volume ∼2500 Å^3^) which is related to insertion of MDH_62-70_ loop, characteristic only for the glycosomal form of MDH. To determine the significance of the identified interaction we tested the binding of Pex5 mutant with a partial deletion of the loop (Pex5_Δ470-480_) with MDH. The Pex5_Δ470-480_ mutant retained the ability to recognize MDH derived PTS1 peptide indicating the deletion had no significant impact on structure of the mutant, though the affinity was slightly (3x) decreased compared to the wild type (Table 1). At the same time the mutant exhibited nearly four times higher affinity for MDH compared to wild type Pex5 (Table 1). This suggests that the Pex5_470-480_ loop sterically clashes with MDH in the distal state of the complex limiting the flexibility and affinity of the interaction. To furhter test the hypothesis, we mutated Pro490 residue which breaks the TPR4 helix at the onset of the Pex5_457-489_ loop into arginine, a residue which we expected to support the expansion of TPR4 helix and thus limit the loop flexibility. The affinity of P490R mutant was compromised more than 3-fold compared to the wild type and the change in affinity was primarily driven by the entropic component, again demonstrating that the intractions at the intersubuint groove of MDH unfavorably affect the complex dynamics and affinity.

### **Pex14**_NTD_ **binding site and noncanonical interactions with Pex5**

We observed weak density beyond the Pex5 TPR domain which was difficult to interpret in any of 3DVA derived maps and was not examined for the purpose of dynamics analysis in prior paragraphs. This density likely corresponded to either a fragment of Pex5 N-terminal domain (NTD) or Pex14_NTD_. To clarify the ambiguity, we reconstituted *in vitro* and solved the structure of the MDH-Pex5 binary complex devoid of Pex14_NTD_ (see methods). The maps for the binary complex revealed no density beyond MDH and Pex5 TPR domain (Supplementary Figure 13), indicating the additional density observed in the ternary complex originates from Pex14_NTD_. In ternary complex map, we improved the density by focused refinement using all collected data. Such data treatment did not account for the dynamics of the MDH/Pex5 interface but resolved the density features in the Pex region and enabled the unambiguous atomic model building of Pex14_NTD_ located on the Pex5 proximal Wxxx(F/Y) motif. The final atomic model of the ternary complex (designated as MP1P) was built in a composite map comprising the consensus MDH density map derived by standard refinement and Pex5-Pex14 density map derived from focused refinement. The model consists of the MDH tetramer, Pex5 residues (327-462, 487-653), and Pex14_NTD_ (residues 21-85) (Figure 4; Supplementary Figure 7).

**Figure 4:**
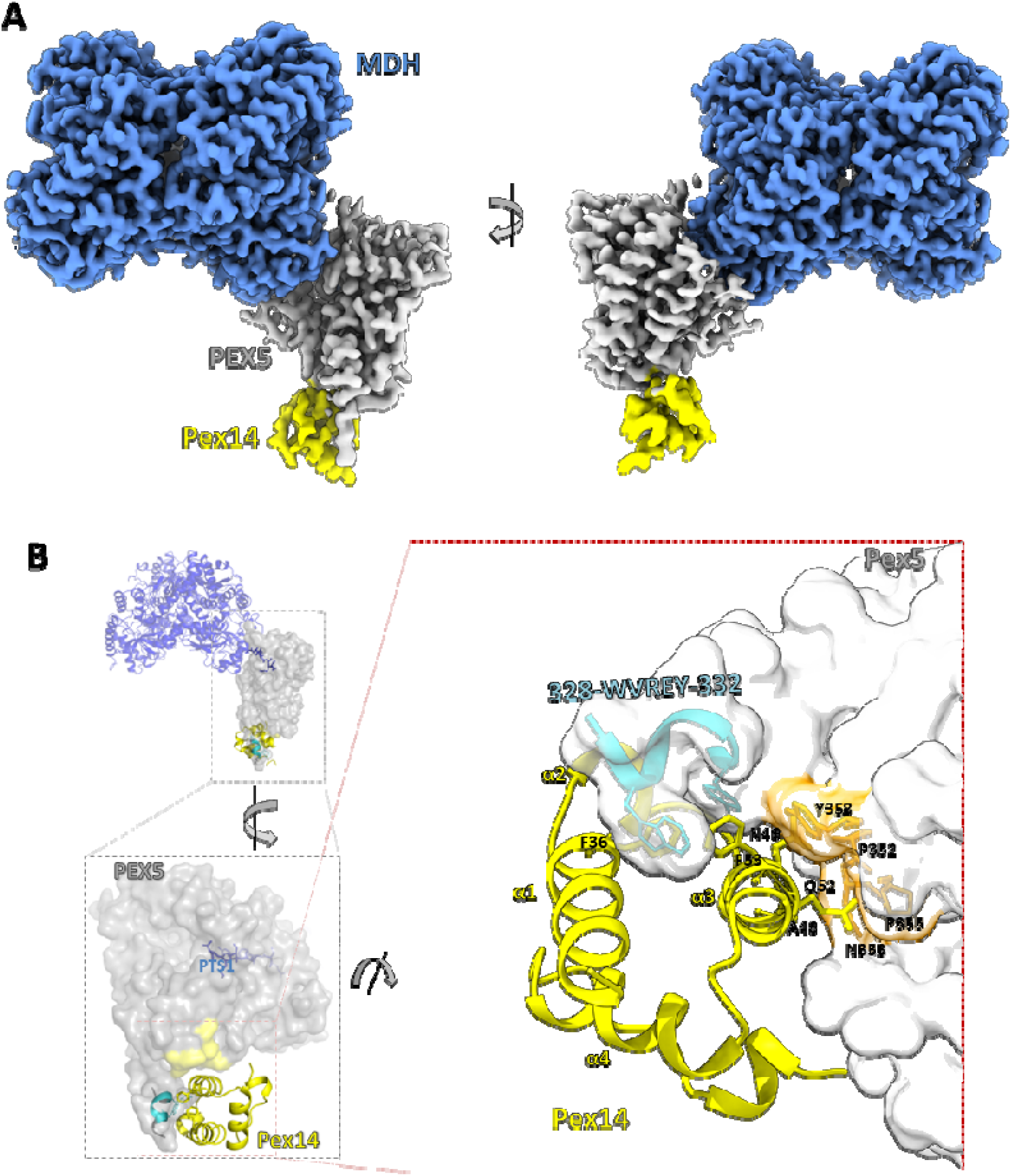
**(A)** Composite cryo-EM map of MDH-Pex5-PEX14_NTD_ ternary complex (MP1P) demonstrating well resolved structural features of MDH, Pex5 TPR domain containing the proximal Wxxx(F/Y) motif (further part of Pex5 NTD is not defined by density suggesting its flexible nature) and Pex14_NTD_. **(B)** Details of Pex14_NTD_ interaction with Pex5 as seen in a model interpreting the map presented in panel A. MDH, Pex5 and Pex14_NTD_ are depicted in blue ribbon, grey surface and yellow ribbon, respectively. Non-canonical interaction region between Pex5 and Pex14_NTD_ along with canonical Wxxx(F/Y) motif mediated interactions are represented by orange and cyan colours, respectively. Helices of Pex14_NTD_ are annotated in yellow. Residues most relevant for the interactions are shown in stick model.

The ternary complex structure constructed in MP1P composite map reveals an extended Pex14_NTD_-Pex5 binding surface involving two distinct interfaces (Figure 4). The canonical interface encompasses Pex5 Wxxx(F/Y) motif^20^ (328-<u>W</u>VRE<u>Y</u>-332) constituting a part of an α-helix (residues 327-345) extending beyond the globular structure of the TPR domain of Pex5 (Figure 4). The core interactions in this region are contributed by bulky sidechains within the motif (underlined) which bury in a hydrophobic groove on the surface of Pex14_NTD_. The canonical interface resembles closely to those observed in crystal structures of short peptides encompassing the Wxxx(F/Y) motif in complex with Pex14_NTD_ from different species^20^. The second, noncanonical interface has not been observed before and involves Pex14_NTD_ helix α3 and 353-NNPYM-358 motif of Pex5 (Figure 4; Supplementary Figures 9, 11). The noncanonical interface contributes ∼800 Å² of buried surface area out of a total ∼1300 Å² comprising the Pex14_NTD_-Pex5 interface. The remaining ∼500 Å² are contributed by the canonical site. The 353-NNPYM-358 region forms an extended interhelix linker separated on both sides from the helical regions by helix breaking P352 and P355. The resolution limits detailed analysis of interactions, but CB atoms are clearly discernable allowing to roughly determine the position of the sidechains. Sidechains of Pex5 N353 and Y358 extend in the direction of Pex14 but seem not to be involved in any specific interactions. The sidechains of A48 and Q52 of Pex14_NTD_ extend in the direction of Pex5, but again no specific interactions can be predicted in the region. Overall, the interface does not resemble protein-protein interaction interfaces, but rather a proximity assembly. To evaluate the contribution of the noncanonical interface to Pex14-Pex5 affinity we designed and tested the affinity of relevant mutants. Affinities were determined in presence of PTS1 peptide to maintain cargo-bound like, closed conformation of the TPR domain of Pex5. The affinity (K_d_) of the wild type Pex14_NTD_ for Pex5_TPR_ domain containing a single proximal Wxxx(F/Y) site, was estimated at ∼53 nM by isothermal titration calorimetry (ITC) (Table 2). The affinity of Pex14_NTD_(Q52A) mutant was identical to the wild type indicating that Q52 does not contribute specific interactions with Pex5. The affinity of Pex14_NTD_(A48F) decreased roughly two times compared to the wild type. This demonstrates that the sidechain of A48 indeed points at the direction of Pex5 as predicted, but the interface is flexible enough to accommodate a bulky sidechain in place of a short sidechain of Ala48 with only insignificant steric hinderance. Mutating Y358 of Pex5 to alanine decreased the affinity of the mutant by roughly 1.5 times compared to the wild type, again as predicted in terms of involvement of Y358 in the interface. This again indicates that the interface is flexible to accommodate significant changes in sidechain properties without large effect on affinity. Pex5(N353E) had almost no effect on affinity, demonstrating that hydrogen bonding is not involved in the interaction. To further substantiate the above findings, we introduced a GSGS-spacer between Pex5_TPR_ domain and Wxxx(F/Y) motif to retain the primary interface, but physically separate the secondary interaction surfaces. The mutation resulted in roughly 3.5-fold decrease in the affinity demonstrating that both the canonical and noncanonical sites contribute to the affinity of Pex14_NTD_ and Pex5, with the canonical interaction having major contribution to affinity.

**Table 2.**
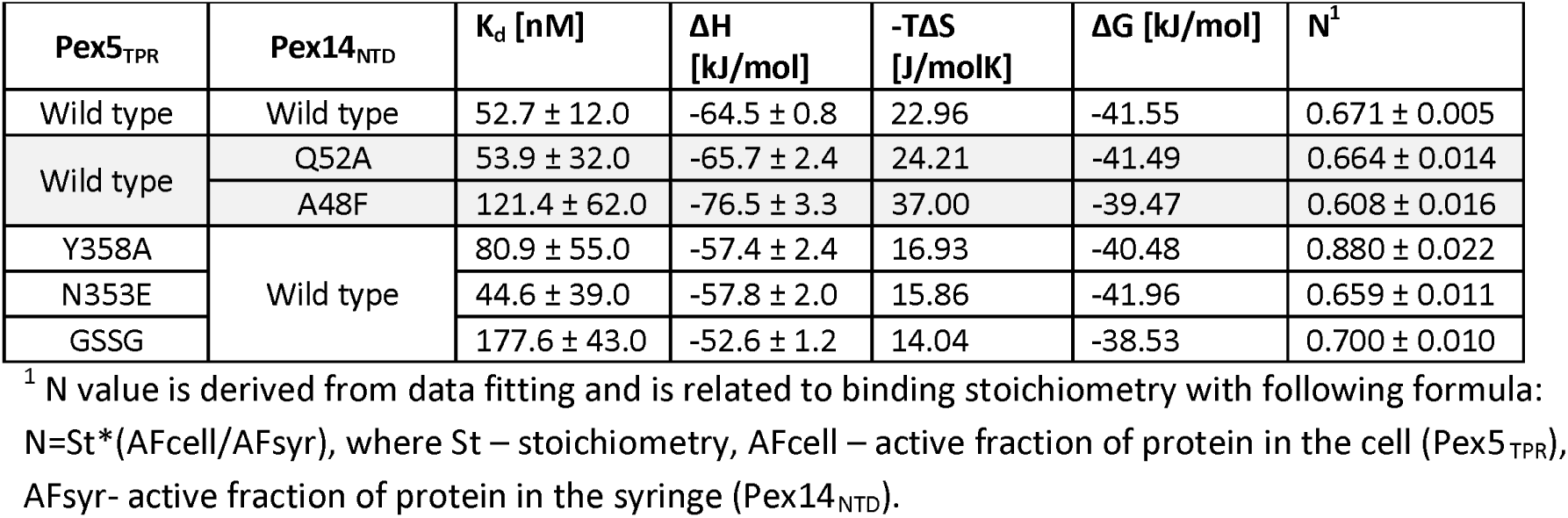
Mutational analysis of Pex14-Pex5 interface. Interaction of indicated mutants was characterized by Isothermal Titration Calorimetry (ITC) in the presence of PTS1 peptide.

## DISCUSSION

Earlier studies have shown that PTS1 guided cargo recognition by Pex5 exhibits both autonomous^11^ (PTS1 mediated, cargo structure independent) and non-autonomous^9^ (cargo structure dependent) characteristics. Non-autonomous model is exemplified by alanine-glyoxylate aminotransferase (AGT) which binding to Pex5 is substantially enhanced by secondary interactions^9^. Conversely, it has been demonstrated that the secondary interface does not significantly contribute to affinity of Pex5 for human sterol carrier protein 2 (SCP2) ^8,11^. Nanomolar affinity for Pex5 of short peptides encompassing PTS1 (in the absence of any secondary interactions) and successful import of PTS1-appended non-peroxisomal proteins into peroxisomes (in the absence of evolutionary optimized secondary interactions) support predominant importance of the autonomous model^33–36^. Our study aligns with the above results by indicating that the secondary interface contributes only weakly to Pex5 affinity for MDH (4 x difference in affinity for Pex5 between WT and GSGS mutant of MDH; Table 1). At the same time, we show that the secondary interactions modulate the dynamic behaviour of Pex5-cargo complex.

Prior studies involving ATG and SCP2 have not accounted for the dynamics of the interface as static crystal structures were analysed^9,11^. Our study demonstrates enhanced affinity of Pex5 for MDH upon abolishing of secondary interfaces (GSGS insertion mutant, Table 1). Similar effect was observed earlier for SCP2^11^. The effects are not pronounced and we postulate that the secondary interactions (as well as the interactions of Pex5_457-489_ loop) only provide proximity clashes which limit the interaction dynamics rather than contribution to affinity. Following our findings in *Trypanosoma*, a comparable dynamic movement of Pex5_TPR_ relative to cargo was identified in yeast^12^. The consensus dynamic component in cargo recognition in two distant organisms suggests that the characteristics may be common to peroxisomal import. Evolutionary adjustment of the interface region seen in peroxisomal MDH in *Trypanosoma* (Supplementary Figure 10) suggests physiological relevance of finetuning the interaction dynamics. We speculate that dynamics plays a role in translocation through peroxisomal membrane and proximity assemblies provide cargo independent mechanism of limiting the dynamics, but further studies are needed to support such hypothesis.

Prior studies thoroughly characterized the interaction of Pex14_NTD_ with short peptides containing the primary recognition motif characterized by consensus sequence of Wxxx(F/Y)^20^, but the binding of Pex14_NTD_ relative to the Pex5 TPR domain remained unclear. Our structure of the ternary complex provides direct visualization of the Pex14_NTD_ binding site in relation to the Pex5 TPR domain. The structure revealed that the Wxxx(F/Y) motif forming the binding site for Pex14_NTD_ is situated within an α-helix which protrudes from the TPR domain, which helix has not been defined in previous crystallographic studies ^7,20^. Such docking of Pex14_NTD_ at Wxxx(F/Y) brings Pex14 in close proximity to TPR4 motif of Pex5. The interface, however, lacks canonical features of protein-protein interactions and rather constitutes a proximity assembly. In support, residues involved in the assembly contribute relatively weakly to affinity as demonstrated by relevant mutants.

There has been ongoing discussion on the role of Pex14 in cargo release^37^. Earlier studies proposed that the binding of Pex14_NTD_ to the Pex5-cargo complex triggers cargo release^38^. In contrast, it has been shown in *Arabidopsis* that the Pex14_NTD_ binding to Pex5 is not sufficient for cargo release^39^. Our complex clearly demonstrates that Pex14_NTD_ binding at the proximal Wxxx(F/Y) site of Pex5 is not sufficient to release MDH cargo. MDH and Pex14_NTD_ bind on opposite sides of Pex5 and do not compete for binding or affect Pex5 structure, which explains why Pex14_NTD_ cannot induce cargo release by binding to proximal Wxxx(F/Y) motif. There are however multiple Pex14 binding sites on Pex5 suggesting that more complicated set of interactions may take place *in vivo*. Additionally, recent literature suggests another mechanism where cargo release is achieved by Pex5 unfolding during recycling through Pex2/10/12 ubiquitin ligase complex^40^ while the binding of Pex14_NTD_ allows extraction of the cargo complex from the translocation pore^13^.

N-terminal domain of Pex5 (Pex5_NTD_) remains intrinsically disordered in solution. Prior low-resolution SAXS study suggested that that Pex5_NTD_ does not undergo significant rigidification upon saturation with Pex14_NTD_, but rather remains in disordered state^41^. Our study supports these earlier conclusions – we have not seen any density accounting for Pex5_NTD_ in any of the maps though full length Pex5 was used for complex reconstitution and amount of Pex14_NTD_ used for reconstitution was equimolar relative to Wxxx(F/Y) sites. These results indicate that at such conditions Pex5_NTD_ remains for the major part intrinsically disordered. This structural observation is interesting in the light of a recent cryo-EM structure of full length Pex14 where a large rod-like coiled coil domain of Pex14 homotrimer is well resolved^42^, while the Pex14_NTD_ domains located on the other side of the peroxisomal membrane remain unsolved. Pex14_NTD_s are positioned in close proximity in Pex14 homotrimer which should facilitate binding to the multiple Wxxx(F/Y) motifs in Pex5_NTD_ thereby increasing avidity. While our structural data unequivocally demonstrates that the binding of single Pex14_NTD_ to the TPR proximal Wxxx(F/Y) motif does not induce structural reorganization of Pex5 NTD, the question remains if the binding of multiple full length Pex14_NTD_s *in vivo* might reorganize Pex5 NTD. Increasing the complexity, recent evidence suggests interactions of TPR distal Wxxx(F/Y) motifs with Pex13 SH3 domain^43^ and with Pex13 YG-motif^13^ domain. It is plausible that these TPR distal Wxxx(F/Y) motifs have evolved to preferentially interact with Pex13 rather than Pex14.

In summary, our findings reveal the dynamic nature of Pex5 binding at MDH and identify noncanonical interfaces complementing the canonical recognition of PTS1 by Pex5_TPR_ and Wxxx(F/Y) by Pex14_NTD_. Mutants within identified interfaces demonstrate their low contribution to affinity and we speculate that the interfaces rather constitute proximity assemblies that limit the conformational dynamics of the interaction to finetune peroxisomal transport.

## MATERIAL AND METHODS

### Protein expression and purification

Genes encoding *Trypanosoma cruzi* MDH, full length Pex5 (residues 1-666, GenBank PBJ69826.1), truncated Pex5 containing a single Wxxx(F/Y) motif (residues 314-666; Pex5_TPR_) and Pex14_NTD_ (residues 21-85) were synthesized *de novo* with N-terminal 6×His, 6×His and 6×His-TEV site tag, respectively. Genes were cloned in pET24(+) vector using NdeI and XhoI restriction sites. Recombinant vectors were transformed individually in *Escherichia coli* BL21 (DE3) for over-expression. Proteins were purified by Ni-affinity (Ni-NTA) and size exclusion chromatography (SEC). Purified Pex14_NTD_ was subjected to TEV protease treatment to remove 6×His-tag. Mutations were introduced by site directed mutagenesis and the mutants were purified as wild type proteins.

### *In vitro* reconstitution and purification of the ternary complex

Purified MDH (tetramer), full length Pex5 and Pex14_NTD_ were mixed at 1:4:12 molar stoichiometry and incubated overnight at 4°C for ternary complex formation. The formed complex was separated from remaining individual components by size exclusion chromatography (SEC) using Superose6 10/300.

### Analysis of the complex formation

The highest molecular weight peak after the void volume of Superose6 10/300 was collected as the potential ternary complex sample and analyzed by SDS-PAGE. 10 µL of sample was mixed with 2x SDS-PAGE sample loading buffer and run on the 4-16% gradient SDS-PAGE gel along with the molecular weight marker in Tris-Glycine running buffer followed by the Coomassie blue staining. The presence of all components was confirmed from the resultant band patterns (Supplementary Figure 1).

### Cryo-EM of MDH - Pex5 - Pex14_NTD_ ternary complex

Cryo-EM data collection: Purified ternary complex was vitrified on holey-carbon grids (Quantifoil R 2/1) using FEI Vitrobot Mark IV. Carbon grids were glow-discharged at 8 mA for 70 sec using LEICA EM ACE200 glow-discharger. Glow-discharged grids were loaded with 3 µL of sample, blotted (blot time: 5 sec; blot force: 5 units) and flash-frozen by plunging into liquid ethane. Frozen grids were screened for the quality of ice and contrast on 200 kV Glacios microscope equipped with Falcon 4 detector prior to data collection on 300 kV Titan Krios G3i Cryo-TEM microscope equipped with Falcon 4 detector at a magnification of ×105k. Each exposure was recorded into 40 dose-fractioned frames with total electron dose of 42.25 e^-^/Å^2^ with a pixel size of 0.86 Å/px. Screening and data collection was performed at cryo-EM facility at the Solaris synchrotron in Krakow, Poland.

Cryo-EM data processing: Data was processed using *cryoSPARC*^44^. Original pixel size 0.86 Å/px was used throughout the processing. Micrographs were motion corrected and subjected to contrast transfer function (CTF) estimation using *patch motion correction* and *patch CTF estimation* jobs, respectively. Protein-particles were picked on a subset of micrographs using *blob-picker*. Picked particles were subjected to template-free 2D classification to generate initial 2D templates. Generated 2D templates were used to pick particles from the whole dataset by *template-picker* with parameters: *particle diameter,* 150 Å; *Lowpass filter to apply* (template / micrograph), 20 Å / 20 Å; *Angular sampling,* 5⁰; *Minimum separation distance,* 75 Å; *Maximum number of picks to consider,* 2000. Bad particles were removed by iterative round of 2D classification. Particles in selected classes representing MP1, MP2 and MP3 complex were used for initial model generation using *ab-initio 3D reconstruction* and subsequent refinement using *homogenous* and *heterogenous refinement* jobs in cryoSPARC. We ended up having the maps for MP1 and MP2 showing satisfactory density for MDH, but not for the Pex region. The 3D-variability analysis (3DVA) ^27^ was used to classify the particles based on heterogeneity. Selected particle clusters and associated volume maps were individually subjected to *non-uniform homogenous refinement* in order to reconstitute individual 3D maps representing different compositional and conformational states of the complex i.e. MP1-c, MP1-d, MP2-c and MP2-d (see Results for definition of states). Local refinement using the local mask on Pex region was performed to improve the map within the Pex region. Local mask was generated to include the density of single copy of Pex region in both distal and close states of MP1-c, MP1-d, MP2-c and MP2-d complexes. All particles, including particles in the class representing MP3, were used for the local refinement using this mask resulting in well resolved density for Pex14_NTD_. Locally refined Pex region map was combined with the consensus map using *Combine map* tool in the Phenix to generate the final map of MP1P complex^45^. Density maps were further sharpened by B-factor based sharpening in cryoSPARC and using EMReady^46^. Overall quality of cryo-EM data and data-processing workflow is depicted in Supplementary Figures 2-7 and Supplementary Table 1.

Model building and refinement: Crystal structure of MDH (PDB id: 7QOZ) and the structure of *T. cruzi* Pex5 TPR domain (PDB id: 7ZT1) were used to fit in MP1 and MP2 maps using *Dock in map* program of Phenix suite^45^. Docked models were subjected to flexible fitting into the map using namdinator web-server^47^. Fitted models were refined against the map by iterative cycles of automatic refinement using phenix real-space refinement^48^ and interactive refinement using COOT^49^ until reasonable geometry and map-to-model correlation were achieved. For MP1P structure, the refined models of MDH and Pex5 from above structures were rigid body fitted in the map, and further refined. The Pex14_NTD_ was then *de novo* built in the map to complete the MDH-Pex5-Pex14_NTD_ model. For Pex14_NTD_ *de novo* model building, its alphafold^50^ predicted structure was docked with in the map. The model was trimmed and refined as needed according the map along with whole complex by COOT^49^, phenix real space refine^45^ and ISOLDE^51^. Validation of model and model *vs*. data for all structures are summarized in Supplementary Figures 3-7 and Supplementary Table 1.

### Cryo-EM structure determination of MDH-Pex5 binary complex and MDH alone

The MDH-Pex5 binary complex was *in vitro* reconstituted similarly as MDH-Pex5-Pex14_NTD_ ternary complex, but without Pex14_NTD._ The cryo-EM structure of MDH alone was solved from a dataset containing MDH-Pex5-Pex14_NTD_ by selection of particles that contained only the MDH tetramer core. The atomic structures were derived as described for the ternary complex. The MDH-Pex5 structure was used to verify the assignment of the density to the Pex14_NTD_ in the MP1P map. The MDH structure was used to verify the assignment of the density to the Pex5_457-489_ loop in the MP1-d and MP2-d maps.

### Data analysis and representation

The analysis, manipulation and representation of cryo-EM maps and 3D structures were performed using COOT^49^, PyMOL (Schrödinger, LLC), UCSF chimera ^52^ and chimeraX ^53^. Sequence alignments were performed and presented using MEGA X software and ESpript server ^54^, respectively.

### Isothermal Titration Calorimetry (ITC)

ITC measurements were performed in 20 mM sodium phosphate buffer containing 100 mM NaCl at pH 6.5 (Pex5-Pex14 interaction) or 100 mM Hepes buffer containing 150 mM NaCl and 2 mM β-mercaptoethanol at pH 7.5 (MDH-Pex5 interaction) using a Nano ITC 2G (TA Instruments). Before the experiments, proteins were extensively dialyzed against the relevant buffer. The binding experiment involved 5 μL injections of Pex14_NTD_ (175 μM) or Pex5_TPR_ solution (250 μM) into the calorimeter cell containing Pex5+PTS1 (20 μM) or MDH (25 μM), respectively. Injections were performed at 300 s intervals. All experiments were conducted at 25 °C with a stirring rate of 250 rpm. The heat of dilution was determined by titration of Pex14_NTD_ or Pex5_TPR_ to protein-free buffer. The data were analyzed using NanoAnalyze software (TA instruments). The N value (defined as N=St*(AFcell/AFsyr), where St reflects stoichiometry [number of binding sites], and AF represent active fractions in cell and syringe), the dissociation constant (K_d_) and the enthalpy change (ΔH) were obtained from fitting the data with binding model. The Gibbs free energy (ΔG) and change in entropy (ΔS) were calculated using −RT lnKd = ΔG = ΔH−TΔS, where R is the gas constant and T is the absolute temperature.

## Data availability

Density maps of complexes, MP1P, MP1-c, MP1-d, MP2-c, MP2-d, MDH-Pex5 and MDH alone are available at the electron microscopy database (EMBD) with accession ids, EMD-40056, EMD-40003, EMD-40008, EMD-40031, EMD-40032, EMD-50340 and EMD-50339 respectively. Refined 3D-structure coordinates of complexes, MP1P, MP1-c, MP1-d, MP2-c, MP2-d, MDH-Pex5 and MDH alone are available at the protein data bank (PDB) with accession ids, 8GI0, 8GGD, 8GGH, 8GH2, 8GH3, 9FEF and 9FEE and, respectively.

## ACKNOWLEDGEMENTS

This work was supported in parts by grants 2017/26/M/NZ1/00797 (construct preparation, crystal structures of selected components) and 2020/39/B/NZ1/01551 (remaining experiments), both to GD, from the Polish National Science Centre. RS was a recipient of NAWA Ulam scholarship. We thank Marcin Jaciuk, Katarzyna Pustelny and Rahul Mehta from Malopolska Centre of Biotechnology of the Jagiellonian University, and Grzegorz Popowicz, Valeria Napolitano and Florian Schlauderer from Helmholtz Center in Munich for valuable discussion, assistance in experiments and providing materials. We acknowledge the access to cryogenic electron microscope at National Synchrotron Radiation Centre SOLARIS supported under contract nr 1/SOL/2021/2 from Polish Ministry of Education and Science. We thank Michal Rawski and Paulina Indyka for assistance. We gratefully acknowledge Polish high-performance computing infrastructure PLGrid (HPC Center: ACK Cyfronet AGH) for providing computer facilities and support within computational grant no. PLG/2022/015912. We thank Klemens Noga from AGH Cyfronet for excellent assistance. We thank Edward H. Egelman (University of Virginia) for making computational facility available for cryo-EM data processing. We thank the MCB structural biology core facility (supported by the TEAM TECH CORE FACILITY/2017-4/6 grant from Foundation for Polish Science) for providing instruments and support. We acknowledge the Core Facility for Crystallography and Biophysics, University of Warsaw (supported by the Foundation for Polish Science under the European Regional Development Fund, TEAM TECH Core Facility POIR.04.04.00-00-31DF/17) for valuable support. The open-access publication of this article was funded by the Priority Research Area BioS under the program “Initiative of Excellence - Research University” at the Jagiellonian University in Krakow.

## CONTRIBUTIONS

RS – devised expression and purification protocols, reconstituted the complex, solved the cryo

EM structure of ternary complex, analysed the data, wrote the manuscript, prepared figures.

OL – solved cryo-EM structures of MDH-Pex5 and MDH alone.

AB – expressed proteins, performed biochemical assays, analysed the results, contributed to writing of parts of the manuscript.

M. J-R. - performed isothermal calorimetry (ITC) and analysed the results.

OL, SP, TS – expressed and purified proteins, performed biochemical assays.

GD – obtained funding, coordinated the project, discussed the results, wrote the manuscript.

