## Supplemental figures and tables for "Structure of Trypanosoma peroxisomal import complex unveils conformational dynamics"

Ravi R. Sonani[1,2], Artur Blat[1,3,*], Małgorzata Jemioła-Rzemińska[4,*], Oskar Lipiński[1,3,$,*], Stuti N. Patel[1,*], Tabassum Sood[1,3,*], Grzegorz Dubin[1,#]

[1] Malopolska Centre of Biotechnology, Jagiellonian University, Krakow, Poland

[2] Department of Biochemistry and Molecular Genetics, University of Virginia School of Medicine, Charlottesville, VA 22903, USA

[3] Doctoral School of Exact and Natural Sciences, Jagiellonian University, Krakow, Poland

[4] Faculty of Biochemistry, Biophysics and Biotechnology, Jagiellonian University, Krakow, Poland

[$] Current Address: Universite Claude Bernard Lyon 1, CNRS, Tissue Biology and Therapeutic Engineering Laboratory (LBTI), UMR 5305, F-69367 Lyon, France

[*]Equal contribution, listed alphabetically

This Supplementary Material file contains total 13 supplementary figures, two supplementary tables and captions of eight supplementary movies.

**Supplementary Figure 1**

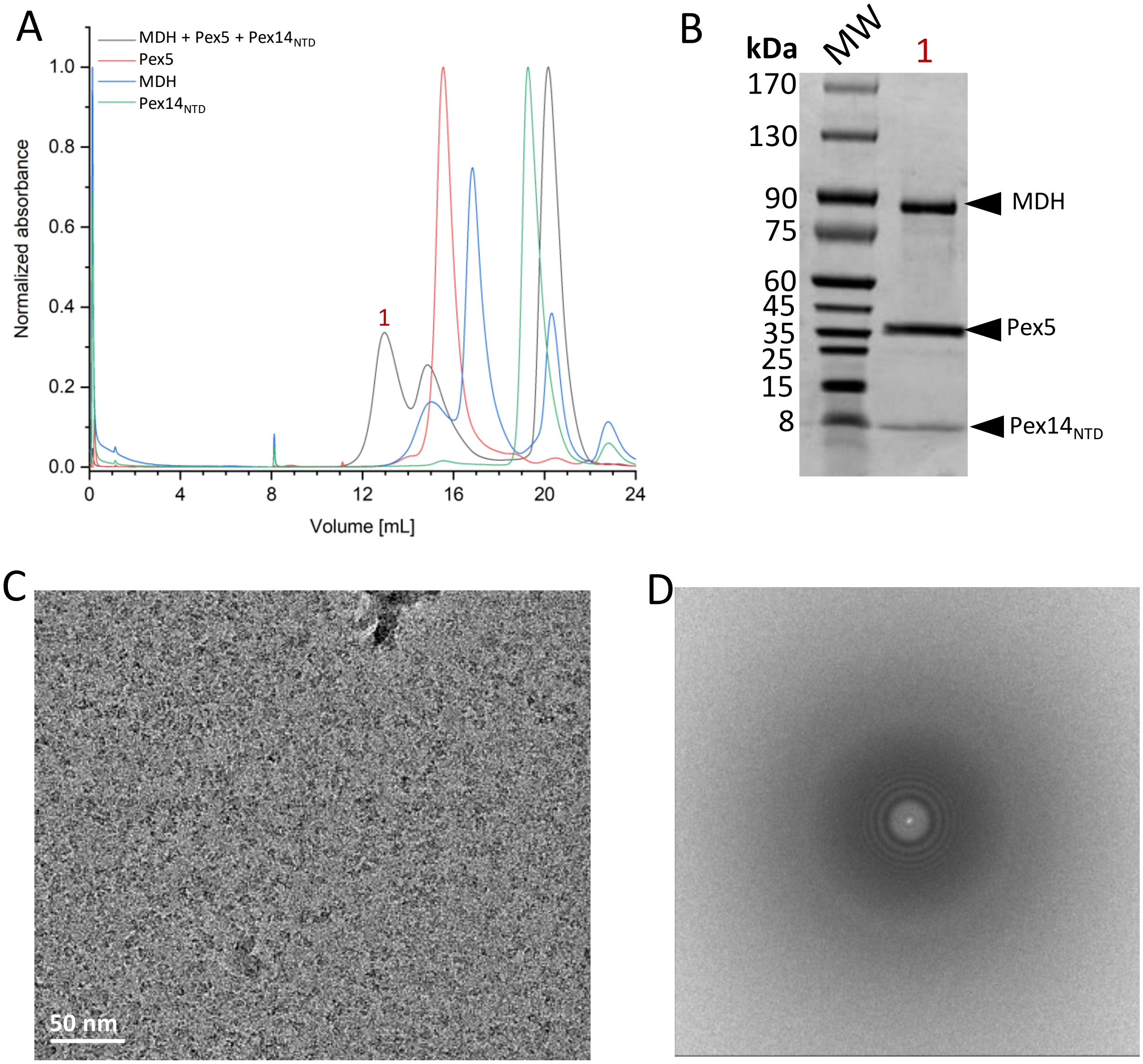


**Supplementary Figure S1**: (**A**) Size exclusion chromatography (Superose6 10/300) profile of *in vitro* reconstituted ternary complex MDH-Pex5-Pex14$_{NTD}$ from *T. cruzi*. MDH, malate dehydrogenase. **(B)** SDS-PAGE of the peak '1' of SEC elution shown in **A**. **(C)** Representative motion-corrected cryo-EM micrograph (defocus range for the whole dataset: -3.0 to -0.9 µm) and **(D)** Power spectrum from the cryo-EM micrograph of ternary complex MDH-Pex5-Pex14$_{NTD}$ shown in **C**.

**Supplementary Figure 2**

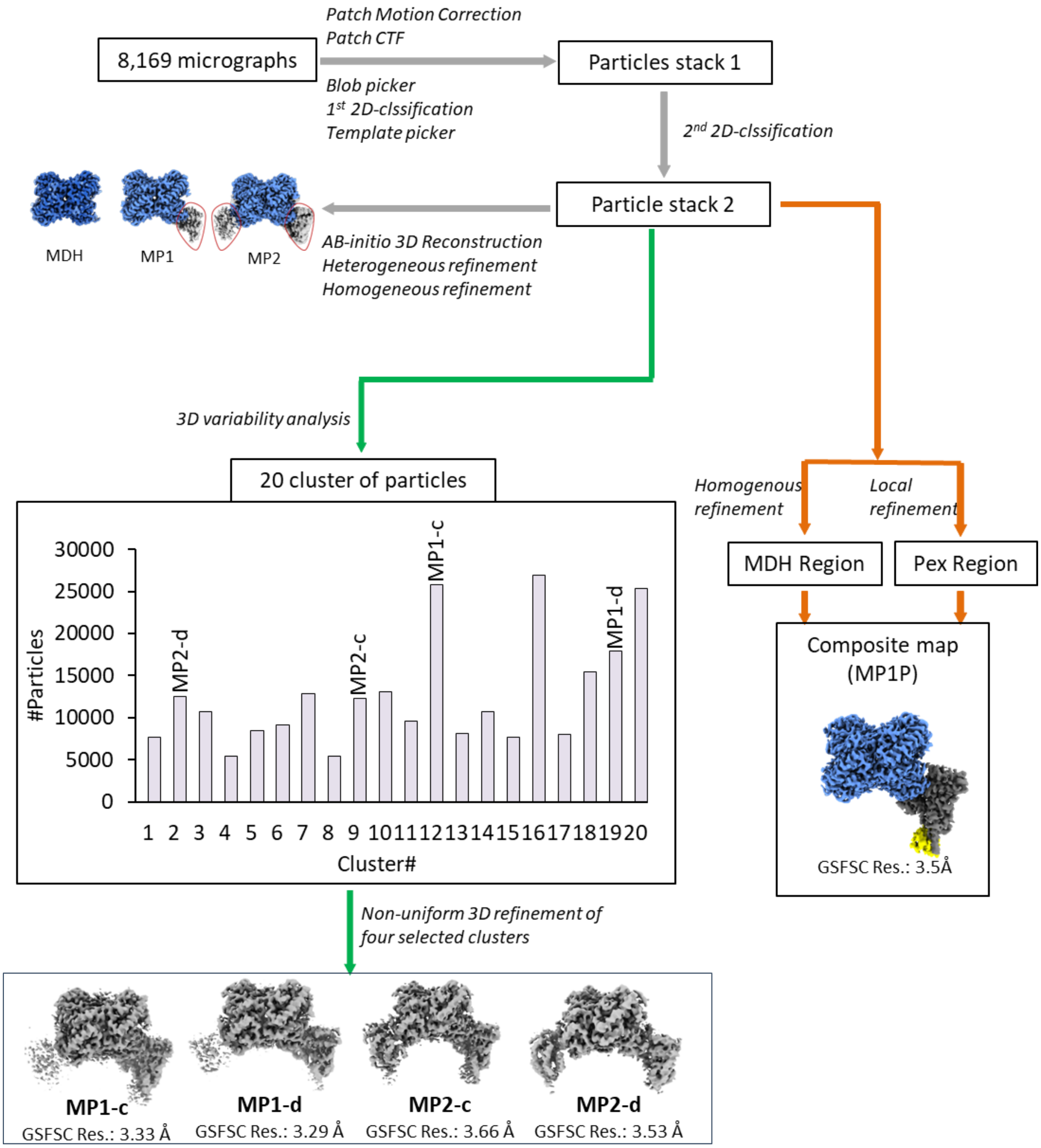


**Supplementary Figure 2**: Cryo-EM data processing workflow used to derive maps of *T. cruzi* MP1-c. MP1-d, MP2-c, MP2-d and MP1P complexes.

**Supplementary Figure 3**

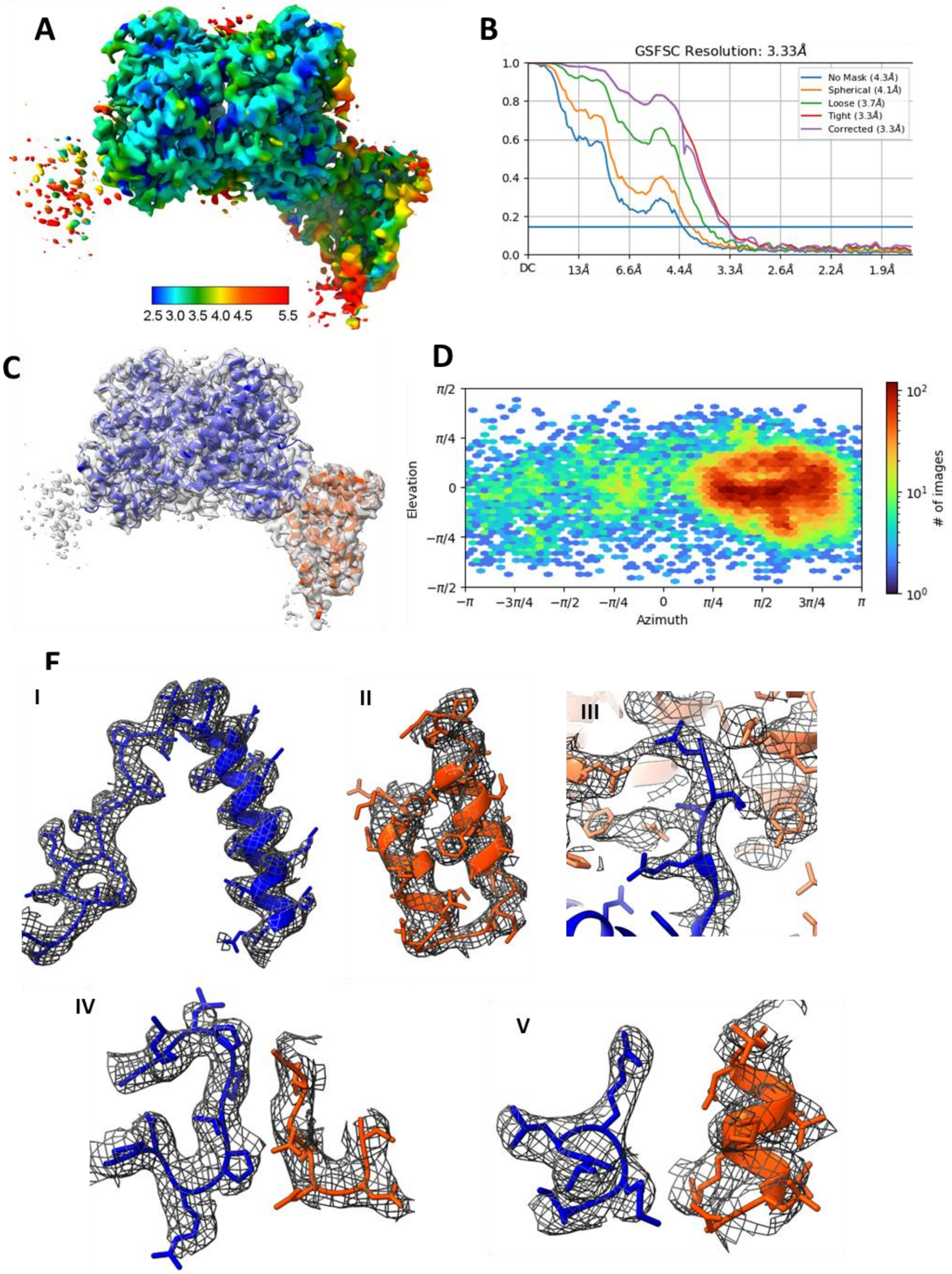


**Supplementary Figure 3**: Quality of cryo-EM structure of *T. cruzi* MP1-c complex. **(A)** Local resolution map, **(B)** Half-maps Fourier Shell Correlation Coefficient (FSC) analysis, **(C)** Overall fit of model into density-map, **(D)** Angular distribution of particles, and **(E)** Model-vs-density fit in representative regions of MDH (I), Pex5 (II), PTS1 (III), MDH-Pex5 interactions (IV and V). MDH and Pex5 are shown in blue and orange-ribbon model, respectively.

**Supplementary Figure 4**

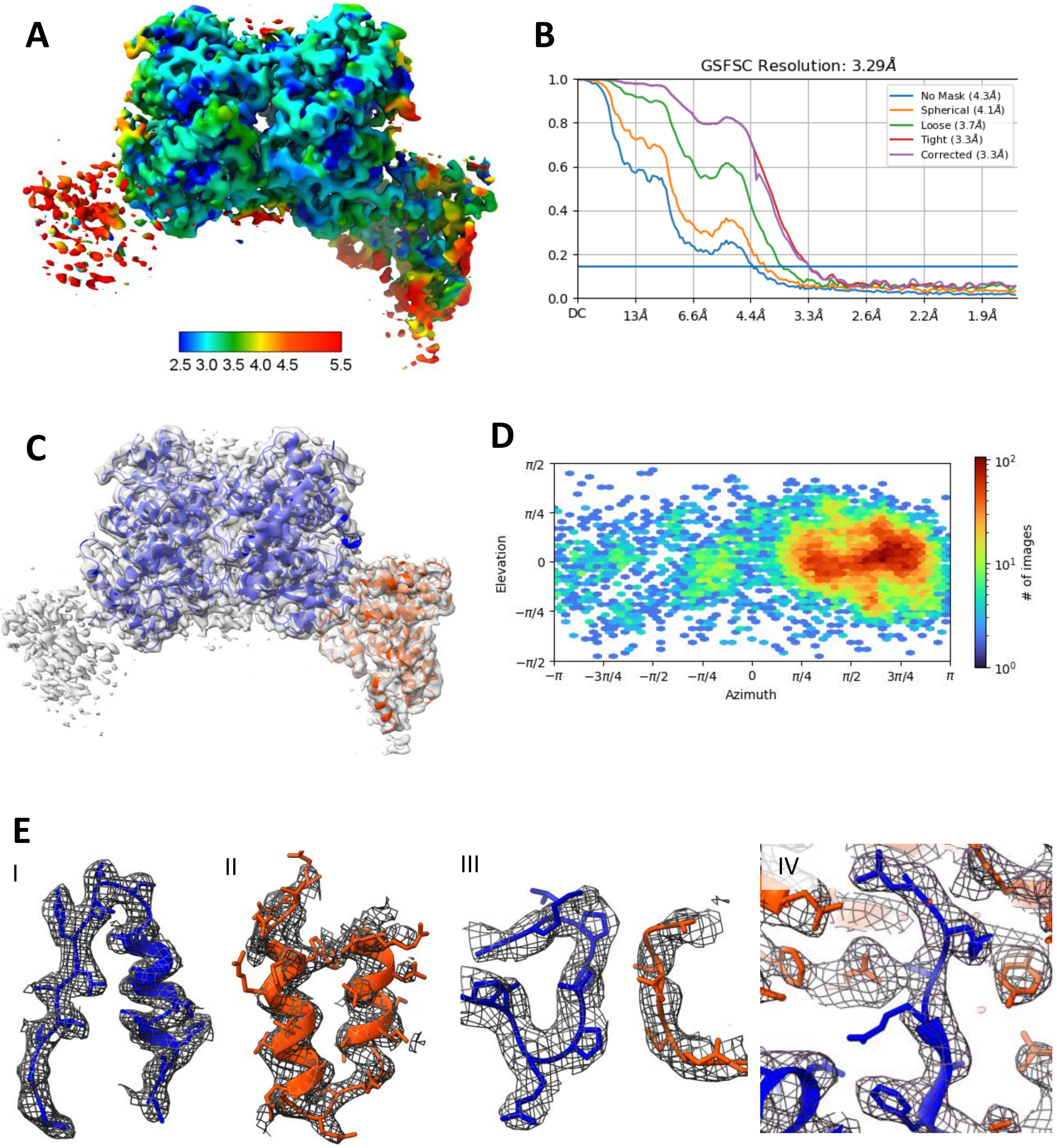


**Supplementary Figure 4**: Quality of cryo-EM structure of *T. cruzi* MP1-d complex. **(A)** Local resolution map, **(B)** Half-maps Fourier Shell Correlation Coefficient (FSC) analysis, **(C)** Overall fit of model into density-map, **(D)** Angular distribution of particles, **(E)** Model-vs-density fit in representative regions of MDH (I), Pex5 (II), MDH-Pex5 interaction (III) and PTS1 (IV). MDH and Pex5 are shown in blue and orange-ribbon model, respectively.

**Supplementary Figure 5**

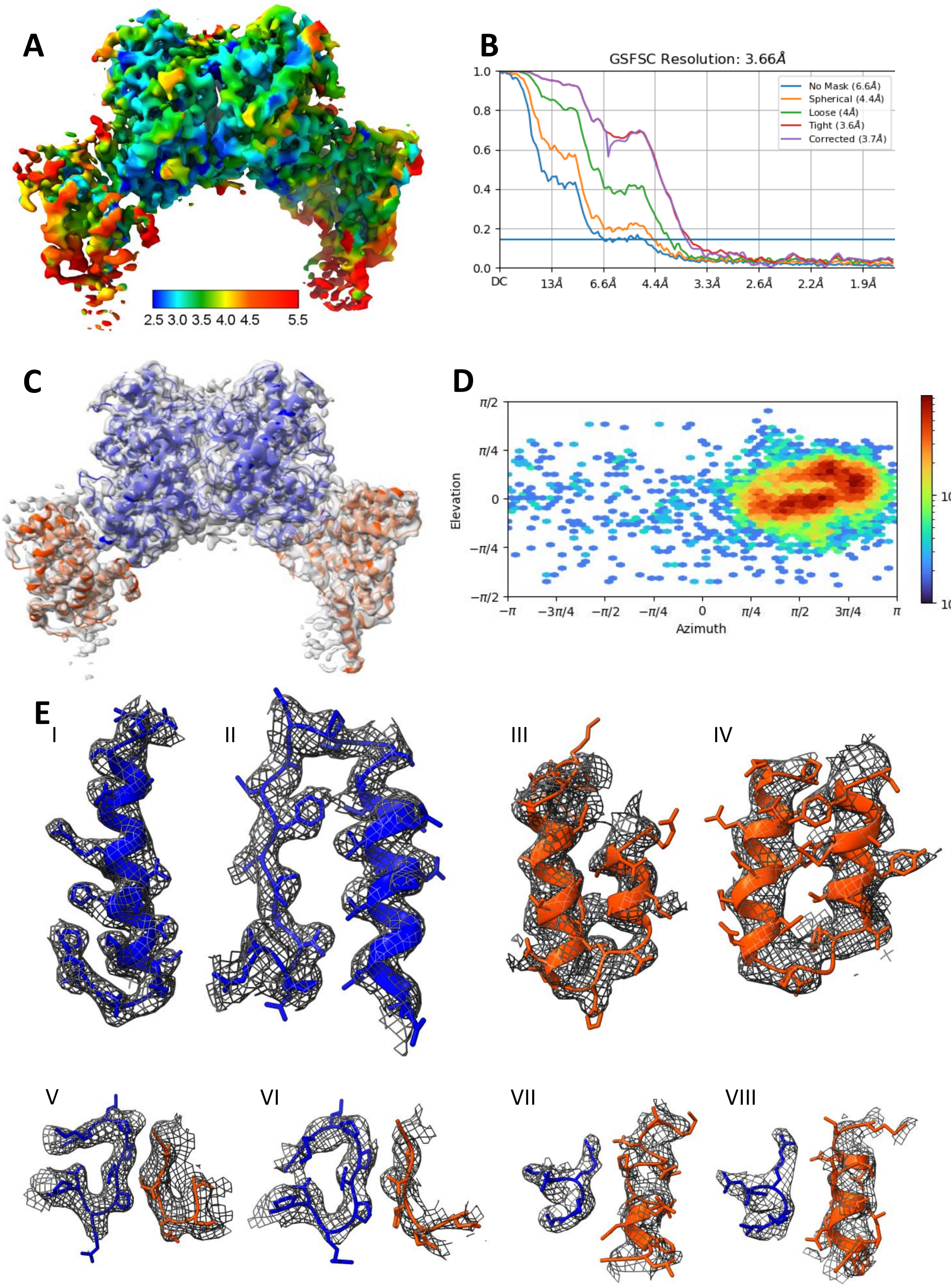


**Supplementary Figure 5**: Quality of cryo-EM structure of *T. cruzi* MP2-c complex. **(A)** Local resolution map, **(B)** Half-maps Fourier Shell Correlation Coefficient (FSC) analysis, **(C)** Overall fit of model into density-map, **(D)** Angular distribution of particles, **(E)** Model-vs-density fit in representative regions of MDH (I, II), Pex5 (III, IV), MDH-Pex5 interaction (V-VIII). MDH and Pex5 are shown in blue and orange-ribbon model, respectively.

**Supplementary Figure 6**

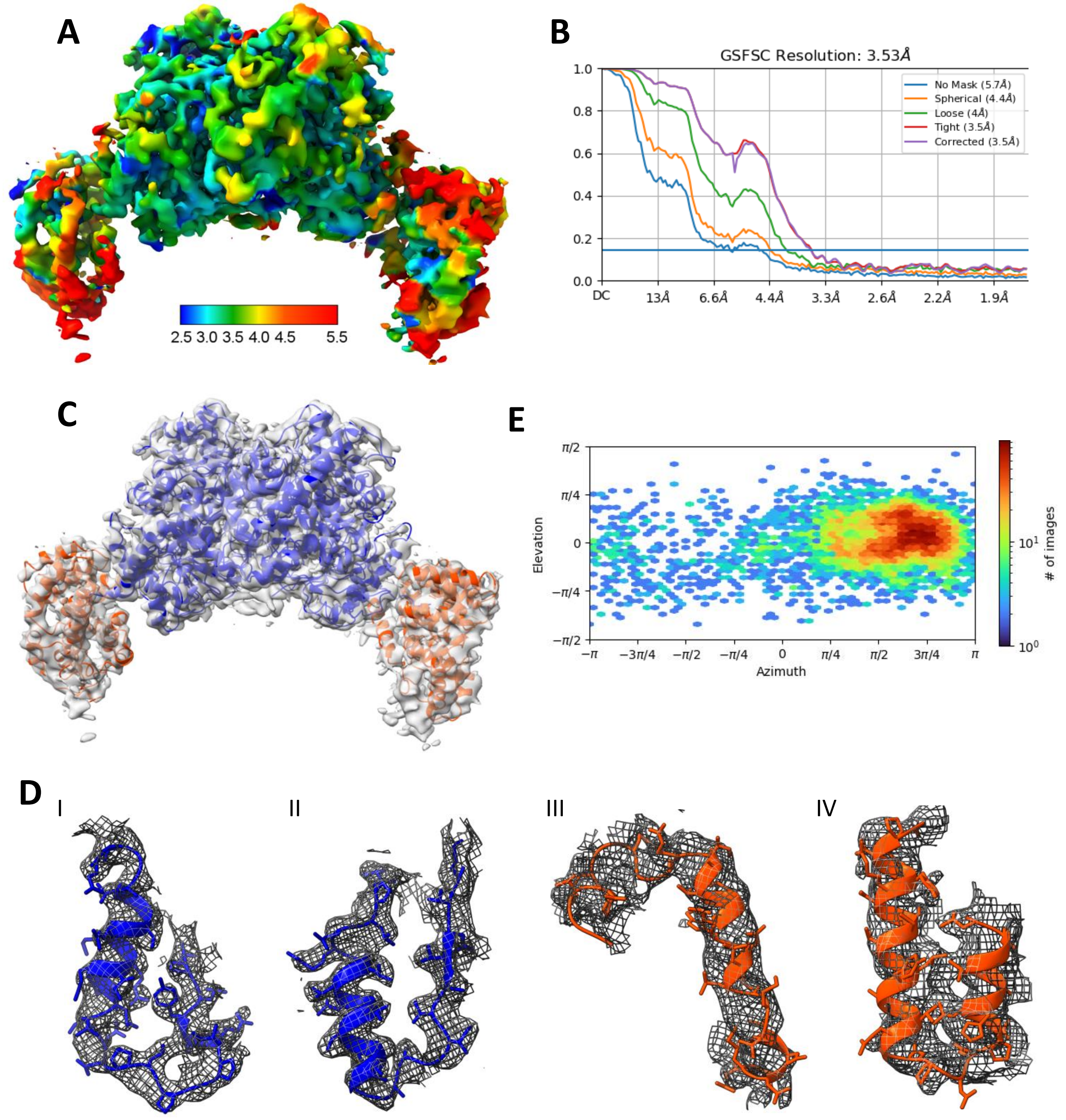


**Supplementary Figure 6**: Quality of cryo-EM structure of *T. cruzi* MP2-d complex. **(A)** Local resolution map, **(B)** Half-maps Fourier Shell Correlation Coefficient (FSC) analysis, **(C)** Overall fit of model into density-map, **(D)** Angular distribution of particles, **(E)** Model-vs-density fit in representative regions of MDH (I, II) and Pex5 (III, IV). MDH and Pex5 are shown in blue and orange-ribbon model, respectively.

**Supplementary Figure 7**

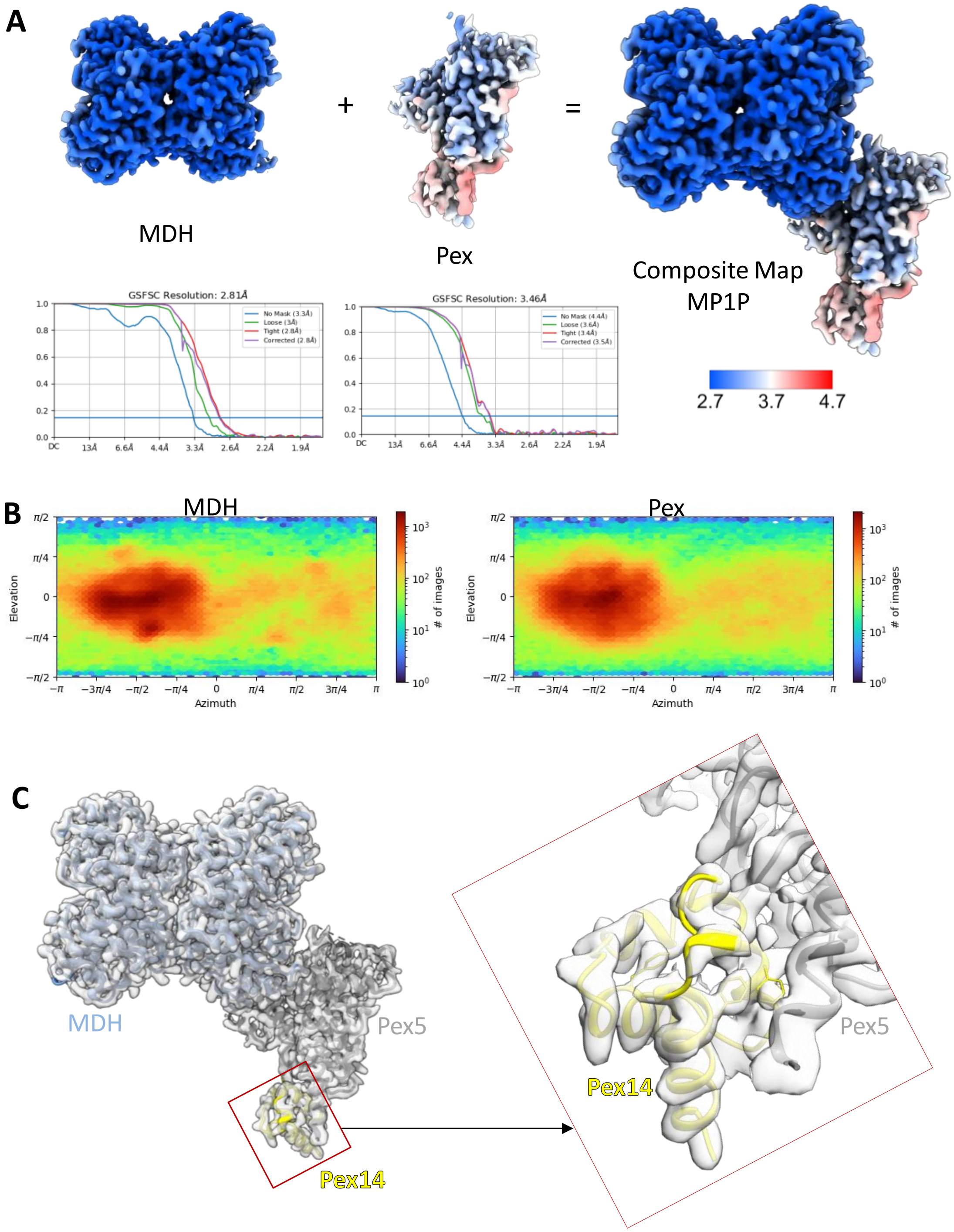


**Supplementary Figure 7**: Cryo-EM structure of *T. cruzi* MP1P complex. **(A)** Individual maps of MDH and Pex region are combined to generate the composite map of MP1P. The local resolution maps and global resolution map-to-map FSC analysis of individual maps are shown. **(B)** Angular distribution of particles, **(C)** Overall model-to-map fit with close up view of Pex14 region.

**Supplementary Figure 8**

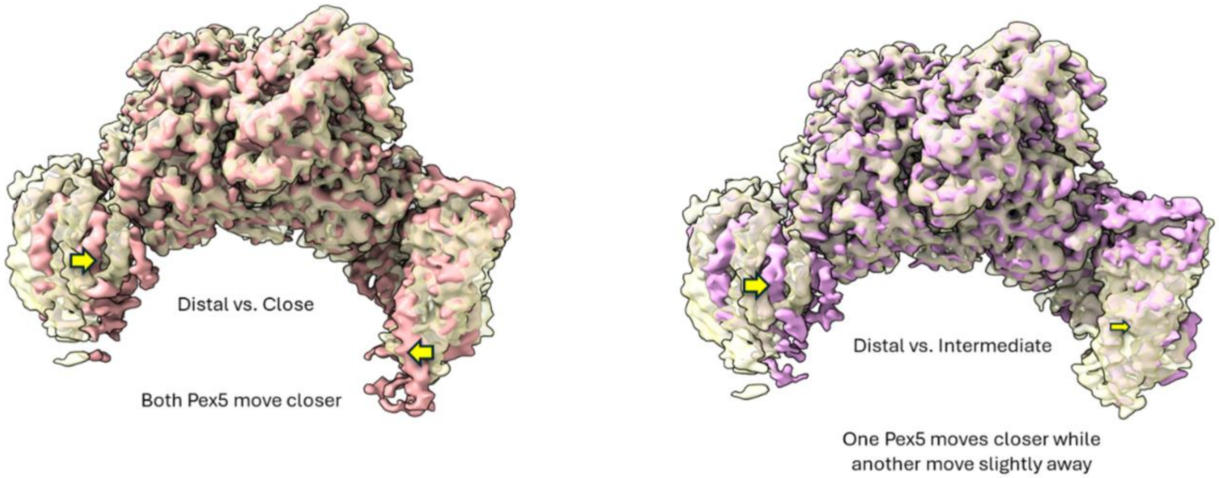


**Supplementary Figure 8**. **Two Pex5 molecules swing with no coordination in MP2 complex.** (Left panel) Overlay of 3DVA derived densities representing distal (gold; largest opening angle of each Pex5 molecule on MDH) and close (pink; smallest opening angle of each Pex5 molecule on MDH) conformations of MP2. (Right panel) Overlay of distal and selected intermediate (violet; small and intermediate opening angle respectively for two Pex5 molecules on MDH) conformations of MP2. Arrows indicate the relative movement of Pex5 component in compared structures. The size of the arrow depicts the extent of the swing motion.

**Supplementary Figure 9**

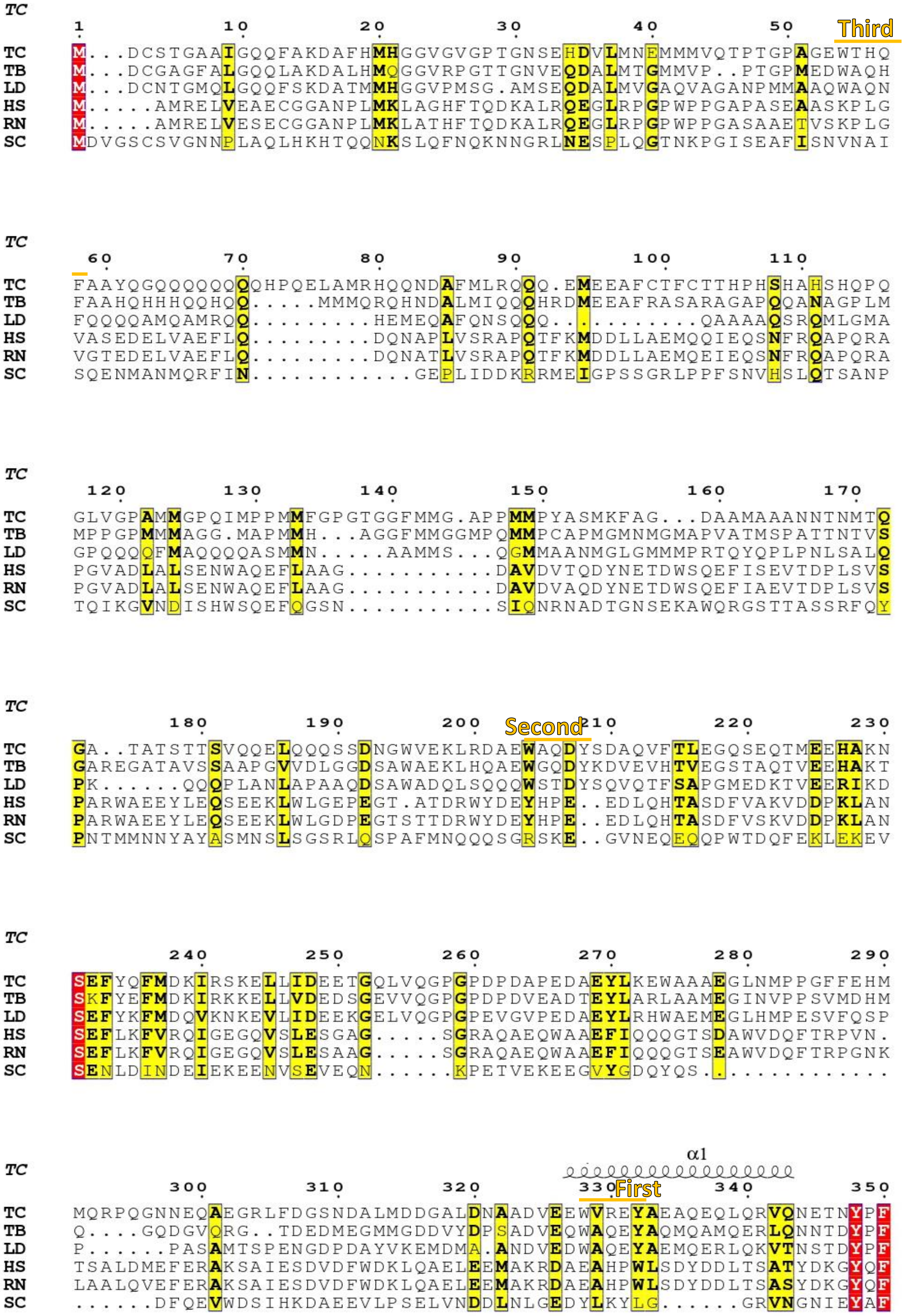


**Supplementary Figure 9**: Multiple sequence alignment of Pex5 homologues from *Trypanosoma cruzi* (TC), *Trypansosoma brucei* (TB), *Leishmania donovani* (LD), human (HS), *Rattus norvegicus* (RN) and *Saccharomyces cerevisiae* (SC). α#, α-helix number. Three Wxxx(F/Y) motifs of in TC Pex5 are designated as First, Second and Third, respectively counting from C- to N-terminus. TPR motifs are indicated by TPR I to TPR VIII. Conserved and partially conserved positions are highlighted by red and yellow, respectively.

**Supplementary Figure 9 (continue)**

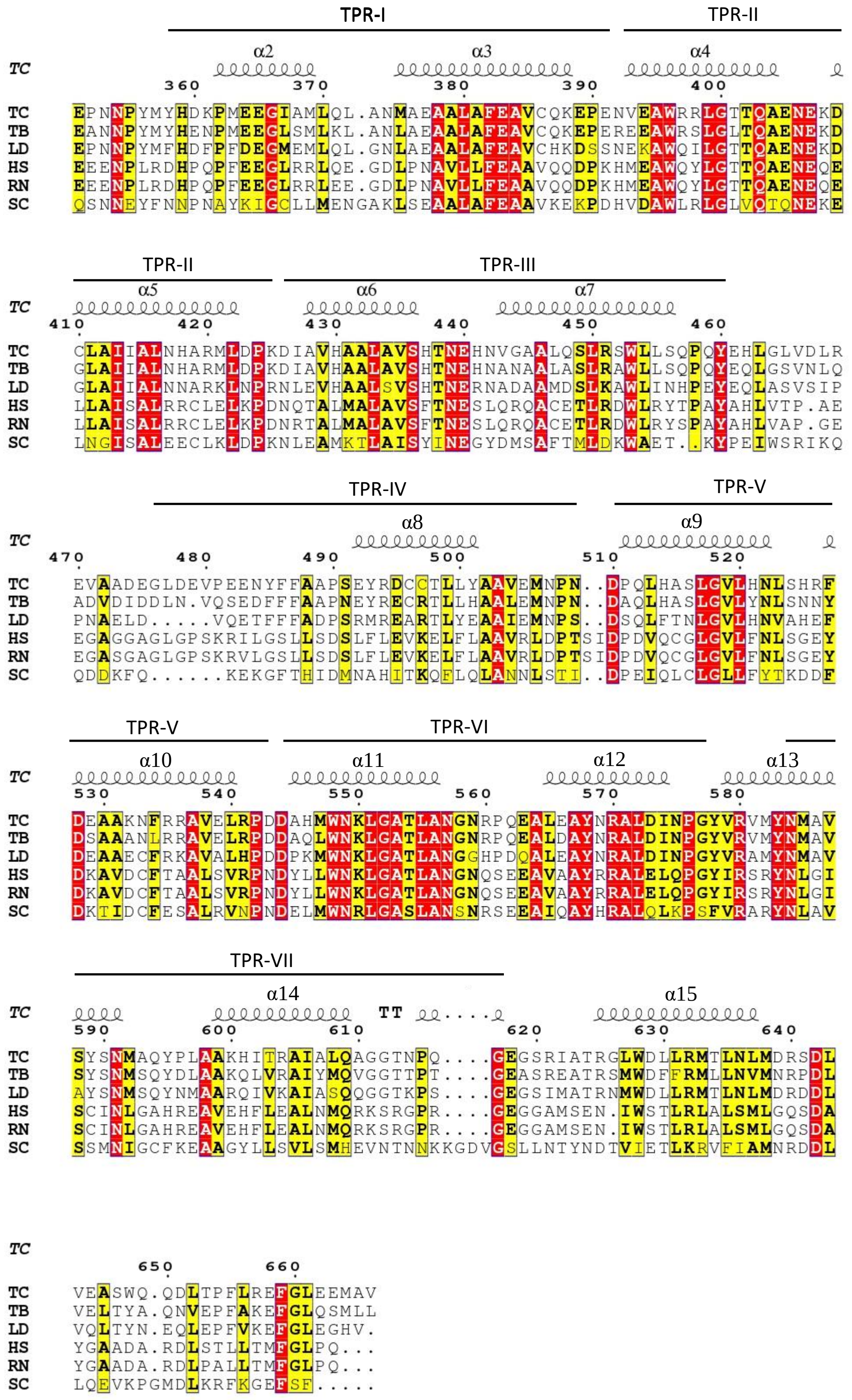

**Supplementary Figure 10**

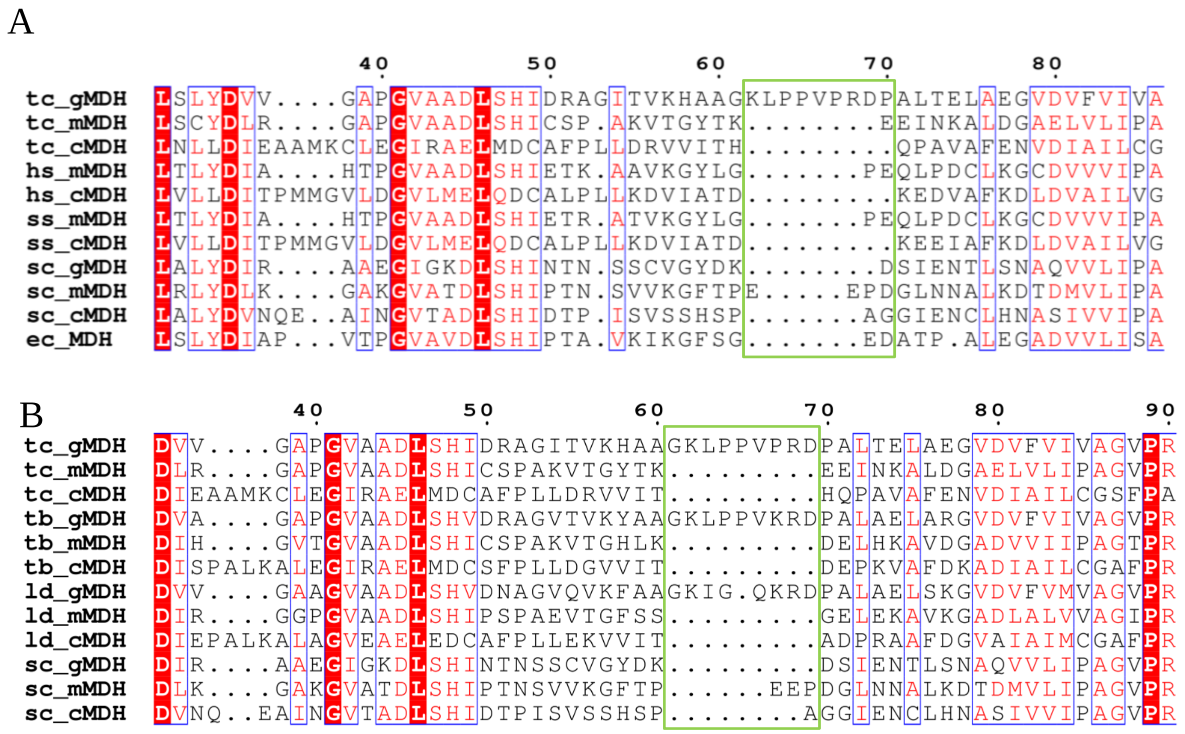


**Supplementary Figure 10**: Multiple sequence alignment of *T. cruzi* glycosomal MDH with subcellular MDH isoforms from different eukaryotes along with the *E.coli* MDH **(A)** and with organisms which compartmentalize glycolytic enzymes in the glycosomes **(B)** highlighting insertion of proline rich 9-mer (62-KLPPVPRDP-70) (green box) in glycosomal MDH. Acronyms: tc, *Trypanosoma cruzi*; hs, *Homo sapiens*; ss, *Sus scrofa*; sc, *Saccharomyces cerevisiae*; ec, *Escherichia coli*; tb, *Trypanosoma brucei*, ld, *Leishmania donovani*; gMDH, glycosomal MDH; cMDH, cytosolic MDH; mMDH, mitochondrial MDH. Conserved and partially conserved positions are highlighted by red and blue boxes, respectively.

**Supplementary Figure 11**

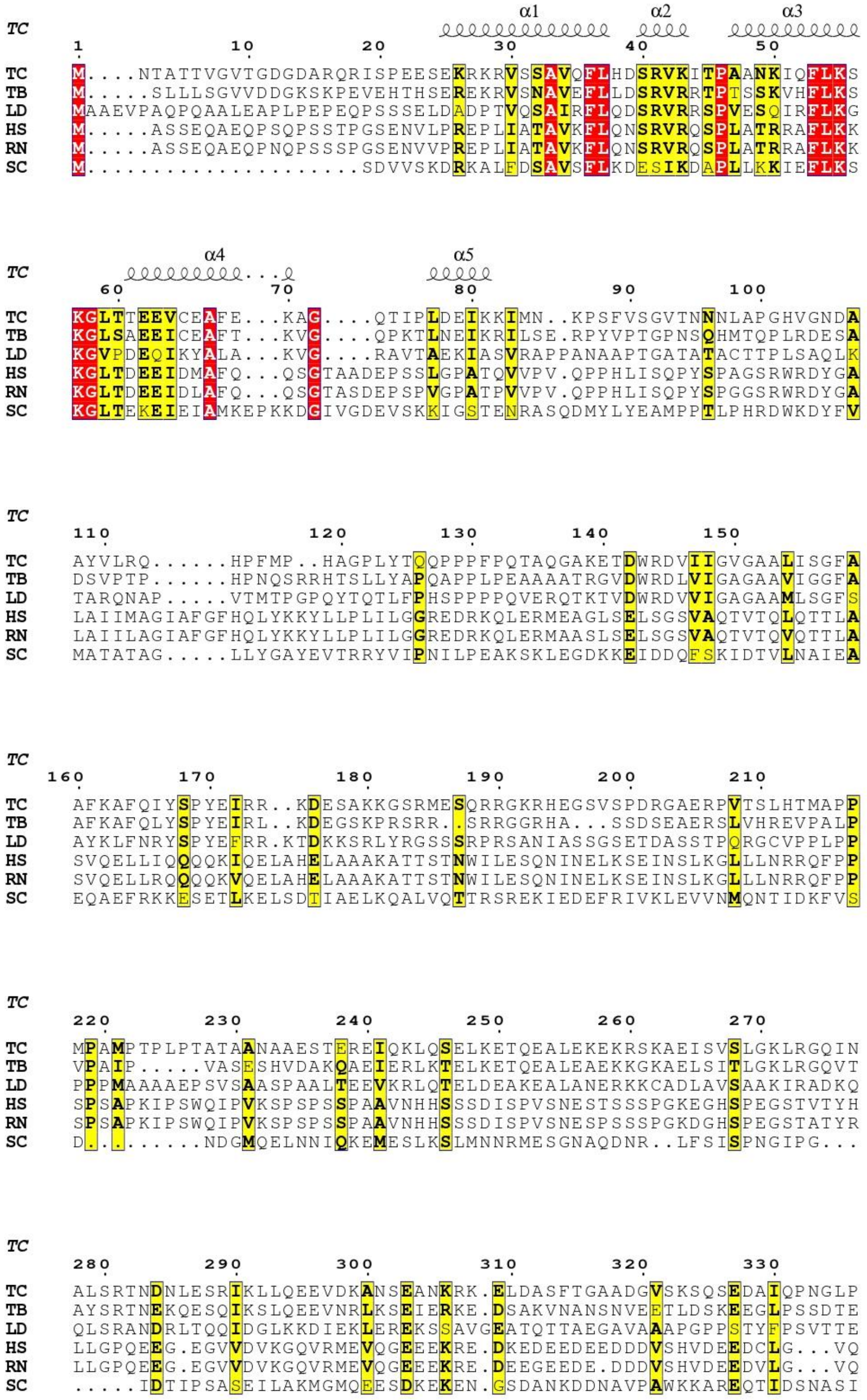


**Supplementary Figure 11**: Multiple sequence alignment of Pex14 homologues from *Trypanosoma cruzi* (TC), *Trypansosoma brucei* (TB), *Leishmania donovani* (LD), human (HS), *Rattus norvegicus* (RN) and *Saccharomyces cerevisiae* (SC). α#, α-helix number. Conserved and partially conserved positions are highlighted by red and yellow color, respectively.

**Supplementary Figure 12**

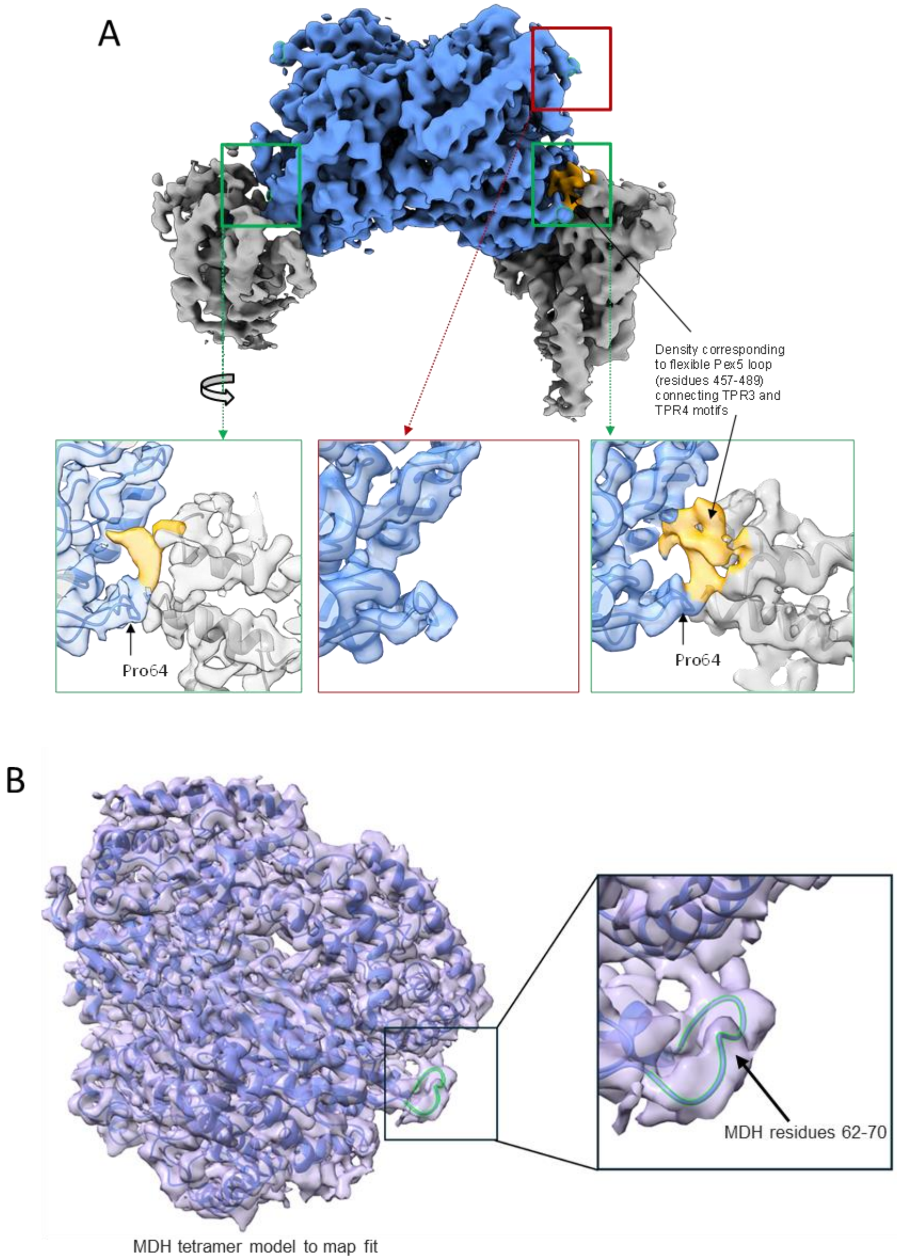


**Supplementary Figure 12:** Non-canonical interactions of Pex5 and MDH in MP2-d complex. **(A)** Density corresponding to the $Pex5_{457\text{-}489}$ loop (gold) transiently interacting with MDH (blue) in MP2-d map. The density is not seen in a groove distal form Pex5 occupied site (red box). See Figure 3 for analogous analysis of MP1-d map. **(B)** MDH model fit in cryo-EM map of MDH tetramer (map was obtained by selecting particles containing MDH only in micrographs used for solution of a ternary complex) . The map shows no density corresponding to the $Pex5_{457\text{-}489}$ loop near the MDH residues 62-70 corresponding to no Pex5 being present in the structure. The presence of discussed density in MP-2 map and absence in MDH map indicates it originates form Pex5 while its positioning indicates it accounts for $Pex5_{457\text{-}489}$ loop.

**Supplementary Figure 13**

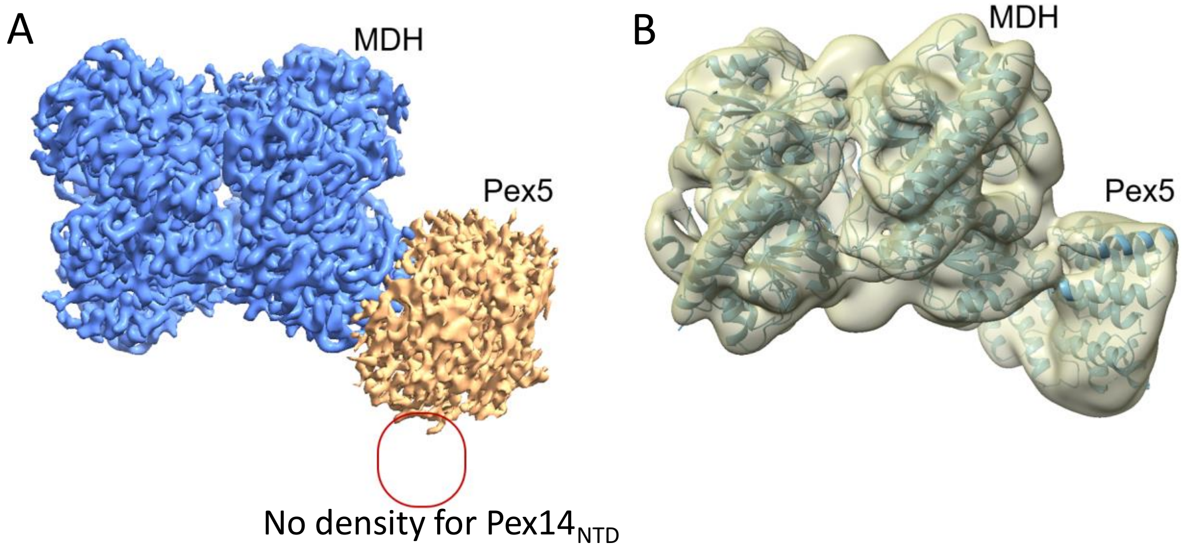


**Supplementary Figure 13**: Cryo-EM map of MDH-Pex5 binary complex obtained in the absence of Pex14. **(A)** High resolution cryo-EM map of MDH-Pex5 showing MDH density (blue) and Pex5 density (brown). Even though full length Pex5 was used for complex reconstitution the map accounts only for TPR domain demonstrating the unstructured, flexible nature of Pex5 NTD. No density is seen in the region indicated by red rectangle corresponding with the fact that complex was reconstituted in the absence of $Pex14_{NTD}$. **(B)** The MDH-Pex5 model fit in low pass filtered cryo-EM map again showing no density corresponding to $Pex14_{NTD}$ corresponding with the fact that $Pex14_{NTD}$ component was not used for complex reconstitution.

**Supplementary Table 1**: Statistics for cryo-EM data collection, image processing and model building

| Data collection and processing | MP1-c | MP1-d | MP2-c | MP2-d | MP1P | MDH alone | MDH-PEX5 |
|---|---|---|---|---|---|---|---|
| **PDB entry** | 8GGD | 8GGH | 8GH2 | 8GH3 | 8GI0 | 9FEE | 9FEF |
| **EMBD entry** | 40003 | 40008 | 40031 | 40032 | 40056 | 50339 | 50340 |
| **Voltage (kV)** | 300 | 300 | 300 | 300 | 300 | 300 | 300 |
| **Magnification** | 105k | 105k | 105k | 105k | 105k | 105k | 105k |
| **Electron exposure ($e^-$/ Å$^2$)** | 42.25 | 42.25 | 42.25 | 42.25 | 42.25 | 42.25 | 42.79 |
| **Defocus range (µm)** | 0.9-3.0 | 09-3.0 | 0.9-3.0 | 0.9-3.0 | 0.9-3.0 | 0.9-3.0 | 0.9-3.38 |
| **Pixel size (Å/pixel)** | 0.86 | 0.86 | 0.86 | 0.86 | 0.86 | 0.86 | 0.84 |
| **Symmetry imposed** | C1 | C1 | C1 | C1 | C1 | D2 | C1 |
| **Particle images (no.)** | 25,845 | 17,940 | 12,276 | 12,554 | 482240 | 312103 | 538505 |
| **Map global resolution (Å)** | 3.33 | 3.29 | 3.66 | 3.53 | 2.8/3.5 | 3.03 | 2.98 |
| **Local resolution FSC threshold** | 0.143 | 0.143 | 0.143 | 0.143 | 0.143 | 0.143 | 0.143 |
| **Refinement** | | | | | | | |
| **Map sharpening (Å$^2$)** | 40 | 35 | 20 | 20 | Sharpened by EMReady | 117.1 | 80.9 |
| ***Model composition*** | | | | | | | |
| **No. of proteins chain (MDH/Pex5/Pex14)** | 4/1/0 | 4/1/0 | 4/2/0 | 4/2/0 | 4/1/1 | 4/0/0 | 4/1/0 |
| **Non-hydrogen atoms** | 11724 | 11798 | 13941 | 13519 | 12191 | 9112 | 11192 |
| **Residues** | 1567 | 1577 | 1846 | 1798 | 1625 | 1236 | 1502 |
| ***Validation*** | | | | | | | |
| **MolProbity score** | 2.30 | 1.93 | 2.08 | 1.96 | 1.39 | 1.73 | 1.75 |
| **Clash score** | 8.47 | 6.89 | 9.08 | 9.60 | 3.56 | 4.99 | 6.46 |
| **Rotamer outliers (%)** | 2.13 | 0.00 | 0.00 | 0.00 | 0.68 | 0.1 | 0.33 |
| ***R.M.D deviations*** | | | | | | | |
| **Bond Length (Å)** | 0.004 | 0.003 | 0.003 | 0.002 | 0.004 | 0.004 | 0.008 |
| **Bond Angle (°)** | 0.724 | 0.640 | 0.706 | 0.672 | 0.687 | 0.582 | 0.796 |
| ***Ramachandran plot (%)*** | | | | | | | |
| **Outliers** | 1.03 | 0.58 | 0.55 | 0.11 | 0.00 | 0.00 | 0.00 |
| **Allowed** | 10.80 | 9.39 | 11.16 | 7.13 | 3.68 | 7.46 | 5.89 |
| **Favoured** | 88.17 | 90.03 | 88.29 | 92.76 | 96.32 | 92.54 | 94.11 |
| **Rama-Z-score** | -1.26 | -0.95 | -1.01 | -0.92 | -0.51 | -1.4 | 0.13 |
| ***Model vs. data*** | | | | | | | |
| **CC (mask)** | 0.83 | 0.85 | 0.81 | 0.80 | 0.83 | 0.87 | 0.87 |
| **CC (volume)** | 0.82 | 0.84 | 0.79 | 0.78 | 0.84 | 0.85 | 0.81 |

**Supplementary Table 2**: The details of cargo, Pex5 and Pex14 variants tested for the ternary complex reconstitution.

| Cargo | Results | Pex5 variants | Pex14 |
|---|---|---|---|
| • Malate dehydrogenase (MDH) (Uniprot: Q4DRD8) | MDH formed ternary complexwhich cryo-EM structure is described in present paper. | • Full length Pex5 (residues 1-666) (GenBank: PBJ69826.1)<br>• Pex5 TPR+ (residue 316-666) (GenBank: PBJ69826.1) | • $Pex14_{NTD}$ (residues 21-85) (GenBank: RNC55913.1) |
| • Phosphofructokinase (PFK) (GenBank: KAF5225211.1) | PFK formed ternary complex as evidenced by SEC, but never gave good 2D class averages in cryo-EM experiment. | | |
| • Glyceraldehyde-3-phosphate dehydrogenase (GAPDH) (UniProt: P22513.1) | GAPDH formed ternary complex as evidenced by SEC but due to small size –of the complex suffered from low signal-to-noise ratio in cryo-EM experiment. No satisfactory 2D class averages were obtained. | | |
| • Phosphoglucose Isomerase (PGI) (PDB: 4QFH) | In our hands PGI did not interact with Pex5. | | |
| • Glycerol Kinase (GK) (GenBank: ESS71147.1) | In our hands GK did not interact with Pex5. The observation was explained by steric occlusion of PTS1 signal in GK dimer as described in[1]. | | |

### CAPTIONS TO SUPPLEMENTARY MOVIES

**Supplementary Movie 1**: Results of 3D variability analysis (3DVA) by cryoSPARC[2] showing the swinging motion of Pex5 relative to the MDH in MP1 complex. The movie is created in UCSF Chimera[3] using the density map series of MP1 generated by cryoSPARC 3DVA[2].

**Supplementary Movie 2**: Morphing between the close and distal conformations of MP1 complex - overview of the complex. The color scheme is the same as in Figure 2. The movie is created in UCSF Chimera[3].

**Supplementary Movie 3**: Morphing between the close and distal conformations of MP1 complex - the view from Pex5 side. The color scheme is the same as in Figure 2. The movie is created in UCSF Chimera[3].

**Supplementary Movie 4**: Results of 3D variability analysis (3DVA) by cryoSPARC[2] showing the swinging motion of Pex5 relative to the MDH in MP2 complex. The movie is created in UCSF Chimera[3] using the density map series of MP2 generated by cryoSPARC 3DVA[2].

**Supplementary Movie 5**: Morphing between the close and distal conformations of MP2 complex - overview of the complex. The color scheme is the same as in Figure 2. The movie is created in UCSF Chimera[3].

**Supplementary Movie 6**: Morphing between the close and distal conformations of MP2 complex - the view from right hand side Pex5 (orientation shown in Supplementary Figure 5 and 6). The color scheme is the same as in Figure 2. The movie is created in UCSF Chimera[3].

**Supplementary Movie 7**: Morphing between the close and distal conformations of MP2 complex - the view from left hand side Pex5 (orientation shown in Supplementary Figure 5 and 6). The color scheme is the same as in Figure 2. The movie is created in UCSF Chimera[3].

**Supplementary Movie 8:** The twisting motion of Pex5 relative to the axis roughly parallel to TPR3 motif. The color scheme is the same as in Figure 2. The movie is created in UCSF Chimera.
